# Discovery of chemical marker for *maidong* (roots of *Ophiopogon japonicus* and *Liriope spicata*): a feature-based molecular networking approach

**DOI:** 10.1101/2024.07.30.605840

**Authors:** F.Y. Lei, L.L. Saldanha, C. Weckerle, L. Bigler

## Abstract

**Background:** Dried tuberous roots of *Ophiopogon japonicus* and *Liriope spicata* are collectively used as *maidong* medicine in China for the same clinical efficacy-nourish *yin* and generate fluids, moisten lung and clear heart fire. Extensive cultivation of these species has necessitated the need for stringent quality control measures. To guide quality control efforts effectively, a comprehensive understanding of metabolomic profiles of *maidong* is essential.

**Methods:** Metabolomic profiling was conducted using ultra-high performance liquid chromatography coupled to a timsTOF Pro hybrid quadrupole-time-of-flight mass spectrometer employing trapped ion mobility spectrometry. Data interpretation was enhanced through feature-based molecular networking (FBMN), uni- and multivariate data analysis (MVDA), and *in silico* annotation.

**Results:** The present study showcases a holistic overview of the metabolomic diversity and variation among *maidong* derived from different origins. Steroidal saponins and homoisoflavonoids were recognized as predominant chemical classes. *Ophiopogon japonicus* predominantly exhibited a variety of homoisoflavonoids, whereas *Liriope spicata* was characterized by a diversity of steroidal saponins. Characteristic metabolites among *maidong* derived from four origins were highlighted. Annotations of 58 metabolites revealed significant inter-species discrimination, with 6 and 36 metabolites critical for regional differentiation in *Liriope spicata* and *Ophiopogon japonicus*, respectively.

**Conclusion:** The current approach effectively discriminated *maidong* from different origins, and facilitated the selection of chemical markers for quality assessment. This approach supports the advancement of quality control strategies for botanical medicines, particularly those derived from multiple origins, ensuring a more rigorous chemical marker selection for botanical medicines.

## 1 Introduction

Dried tuberous roots of *Ophiopogon japonicus* (Thunb.) Ker Gawl together with *Liriope spicata* Lour. (as substitute), both belonging to Asparagaceae, are collectively referred as *maidong* medicine in Traditional Chinese medicine (TCM) (Chinese Pharmacopoeia Commission, 2020). For millennia, *maidong* has been consumed as a monotherapy to nourish *yin* and generate fluids, moisten lung and clear heart fire (Chinese Pharmacopoeia Commission, 2020; Lei et al., 2021). Meanwhile, it is extensively employed in various decoction prescriptions. For instance, *Shengmai San*, a traditional prescription, consists of *maidong* (*O. japonicus*), ginseng (*Panax ginseng* C.A.Mey) and schisandra berry (*Schisandra chinensis* (Turcz. Baill) (Ouyang et al., 2022). In addition, a modern derivative of *Shengmai San*-*Shenmai* injection has been developed and used to treat diverse cardiovascular diseases (Ouyang et al., 2022).

The extensive cultivation of *maidong* in China has led to significant variation in quality, necessitating stringent control measures (Lei et al., 2024). Accordingly, several (inter-)national pharmacopoeias and monographs have established quality assessment standards. Typically, chemical markers are selected via a hypothesis-driven approach, where compounds previously identified as responsible for clinical efficacy are chosen for targeted quantification (Wolfender et al., 2013). For instance, Chinese Pharmacopoeia mandates the evaluation of total saponin content for quality assessment. Ophiopogon D and methylophiopogonanone A (MONA) are regulated in Hongkong Chinese Materia Medica Standards and European Pharmacopeia, respectively (Council of Europe, 2023; Hong Kong Department of Health, 2010). Additionally, other compounds such as methylophiopogonanone B, Ophiopogonanone E, and Ophiopogonanone D have been explored to elucidate the quality variation in *maidong* (Jiang et al., 2022; Lin et al., 2010; Sun et al., 2020; Tan et al., 2019). Nevertheless, the comprehensive array of *maidong* metabolites and their respective pharmacological properties remain insufficiently understood, leading to the selection of markers from a non-holistic perspective. Consequently, an untargeted metabolomics approach is required for achieving a thorough understanding following a hypothesis-generating manner (Aksenov et al., 2017; Houriet et al., 2020; Wolfender et al., 2013).

The handling of untargeted metabolomic data, while challenging, has been considerably advanced by the rapid development of analytical instruments with enhanced sensitivity and resolution, e.g., high resolution mass spectrometry for efficient and accurate data collection. This advancement is further complemented by advanced platforms and statistical analysis tools, e.g., Global Natural Products Society (GNPS, https://gnps.ucsd.edu), feature-based molecular network (FBMN) (Nothias et al., 2020), multivariate data analysis (MVDA) (https://www.metaboanalyst.ca/), *in silico* structure annotation tools (da Silva et al., 2018; Wandy et al., 2018).

The present study was designed to obtain a comprehensive overview of 1) the metabolomic diversity of *maidong* derived from different origins, 2) inter-species level variation between *Ophiopogon japonicus* and *Liriope spicata*, 3) intra-species variation between different geographical origins, and to 4) uncover characteristic chemical markers. Data was collected following an untargeted metabolomics experiment approach using ultra-high performance liquid chromatography coupled to a timsTOF Pro hybrid quadrupole-time-of-flight mass spectrometer employing trapped ion mobility spectrometry (UHPLC-timsTOF). Feature-based molecular network (FBMN) was performed to organize the MS data and highlight structure related compounds (Nothias et al., 2020). This network optimizes the utilization on MS1 and MS2 data, incorporating quantitative metrics for robust downstream metabolomics statistical analysis (Hou et al., 2019). Moreover, integration of FBMN, uni- and multi-variate analysis has facilitated data interpretation. The compounds in the MN were annotated following a taxonomic informed approach using *in silico* spectral database (Allard et al., 2016; Rutz et al., 2019). Overall, our comprehensive MS-based approach not only offers an overview of *maidong*’s metabolomic diversity but also identifies characteristic metabolites crucial for quality control. This method allows a generic and automated identification of chemical markers in phytochemically similar species, particularly medicinal plants sourced from multiple species and locations, addressing a critical gap in current TCM quality control.

## 2 Material and method

### Plant material

For metabolomics analysis of *maidong* from different regions, tuberous roots of *Ophiopogon japonicus* and *Liriope spicata* were collected in production fields located at Sichuan (31.4567, 102.8430), Zhejiang (27.9942, 120.6993), Hubei (31.1678, 112.5881) and Shandong (36.0668, 120.3826) provinces (in yellow color) in China. Samples were collected between February and June in 2021 (Fig. 1, in yellow color). In total, samples from 21 individuals for *Ophiopogon japonicus* and 21 of *Liriope spicata* were obtained. The details for each region sampled, together with the commercial samples are available in the Table S1 and Table S2.

**Fig. 1.**
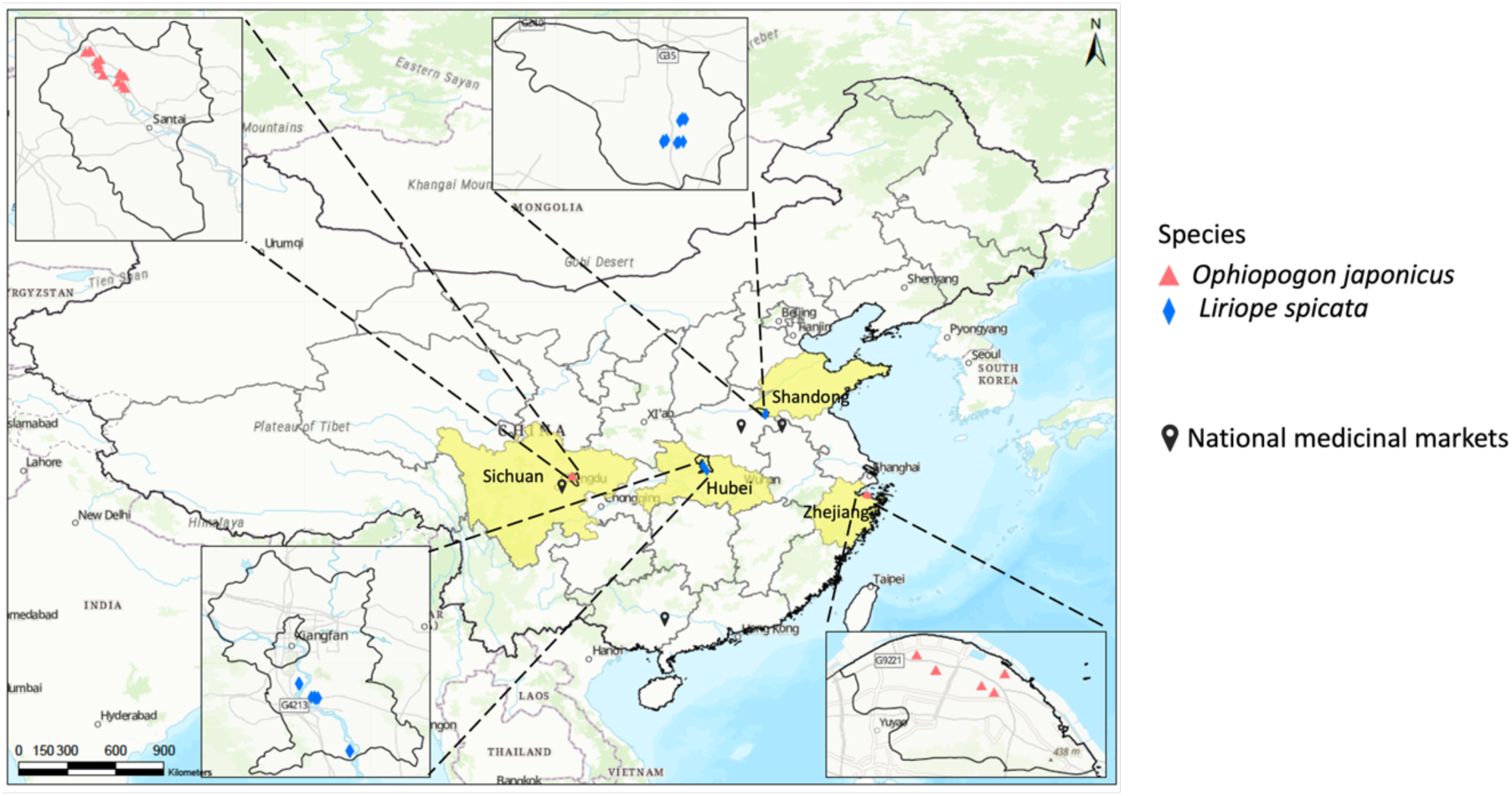
Geographical distribution of four regions where *maidong* is produced in China (Red triangle and blue diamond represent the sites where specimens of *O. japonicus* and *L. spicata* were collected in the present study, respectively).

Tuberous roots were separated from the rest of the plants and rinsed with water. They were dried at 60 °C in the oven and milled using Qiagen Retsch TissueLyser II (20 mm stainless steel ball; 30 Hz frequency; 2 min). The dried and powdered roots were stored with silica gel for later extraction.

For metabolomic analysis of commercial *maidong* samples, tuberous material (34 samples in total) was purchased in national medicinal markets (in Anguo, Yuzhou, Yulin and Chengdu; batch list see Table S2) between February and June in 2021. All samples were milled using Qiagen Retsch TissueLyser II (20 mm stainless steel ball; 30 Hz frequency; 2 min) and stored with silica gel for later extraction.

### Metabolite extraction

Aliquots of 100 mg of dried and powdered roots were extracted with 1 mL 80% MeOH (*v/v*, MeOH/H_2_O). Ampicillin C_16_H_19_N_3_O_4_S [M+H]^+^ ion (*m/z* 349.1096) was used as an internal standard and added into the extraction solvent (80% MeOH). The samples were sonicated for 60 min at room temperature and double centrifuged (14,000 rpm; 4 °C; 15 min). The supernatants were collected into 1.5 mL vials and stored at −20 °C prior to metabolomic profiling.

### LC-HRMS Analysis

Liquid chromatography was performed on a Vanquish Horizon UHPLC system by *Thermo Fisher* (Waltham, USA) connected to a timsTOF Pro mass spectrometer (Bruker Daltonics, Bremen, Germany). Chromatographic separation was achieved at 40 °C on an Acquity HSS T3 column (100 × 2.1 mm, 1.8 µm; *Waters*, Milford, MA, USA). The mobile phase composition was: solvent (A) water with 0.1% formic acid and (B) acetonitrile with 0.1% formic acid. The following gradient was applied at a constant flow rate of 0.5 mL/min: (i) 5% B from 0.0-1.0 min; (ii) linear increase to 100% B until 10 min; (iii) holding 100% B until 13.0 min; (vi) back to the starting condition of 5% B until 13.1 min; and equilibration for 1.9 min until the next run.

Mass spectrometry (MS) data were acquired in both positive and negative ionization mode at 4500 V capillary voltage and 500 V endplate offset with a nitrogen nebulizer pressure of 2.2 bar and dry gas flow of 10 L/min at 220°C. Spectra were acquired at 12 Hz scan rate in the mass range from *m/z* 50 to 1,300 at 40,000 resolution (*m/z* 498 full width at half maximum).

Before running a sequence, the mass analyzer was calibrated using ESI-L Low Concentration Tuning Mix (Agilent, USA), a mass deviation below 2 ppm was tolerated. The mass analyzer was further calibrated between *m/z* 90 and 1246 at the beginning of each LC run with sodium formate solution that was injected using a 6-port-valve with a 20 μL loop at a MS resolution of ca. 46,000 (@ *m/z* 634). Dynamic MS/MS acquisitions were performed in a data-dependent mode at a collision energy of 20-30 eV, 2 to 8 *m/z* isolation width, at 16 to 30 Hz scan rate in a mass range between *m/z* 50 and 1,300, (dynamic MS/MS acquisition), and with a maximum injection time of 50 ms. N_2_ was used as a collision gas.

Plant extracts were injected in a randomized sequence and systematically intercalated at regular intervals with blank and quality control (QC) samples. The quality control samples were prepared by pooling 20 µL aliquots from each extract.

### LC-HRMS data processing

The raw data were converted to mzXML using the ms-convert platform tool-ProteoWizard (Adusumilli and Mallick, 2017) and processed in MZmine (vers. 3.4.16) following protocol (Heuckeroth et al., 2024). The following parameters were used for data processing: noise level at 1.0 × 10^3^ for MS1 and 0 for MS2. The ADAP chromatogram builder was set as: minimum consecutive scans of 5, minimum intensity for consecutive scans of 1.0 × 10^3^, minimum absolute height of 7.0 × 10^3^, and *m/z* tolerance of 5 mDa (or 10 ppm). Parameters for ADAP feature resolver were selected as follows: *m/z* and RT range for MS2 scan pairing were 10 mDa and 0.2 min, S/N threshold of 90, minimum feature height of 1.0 × 10^3^, coefficient /area threshold of 50, peak duration range of 0.05-1.5 min, and the RT wavelet range of 0.02-0.05. Raw data was preprocessed with *m/z* tolerance of 5 mDa (or 10.0 ppm) and an absolute RT tolerance of 0.01 min to get rid of isotopic distributions and molecular ion adducts. The chromatograms were then aligned by applying join aligner, with *m/z* tolerance of 5 mDa, absolute RT tolerance at 0.05 min, and weights for *m/z* and RT at 1. The missing feature list was then completed by a gap filling approach of the same RT and *m/z* range gap filler module, with *m/z* tolerance of 0.005 Da. The feature list was filtered by row filter module with minimum aligned features of 6. Furthermore, only features with MS2 spectra were kept for further analysis. Eventually, a feature list with 1126 features having an associated MS2 spectra was obtained. This feature list was exported for statistical analysis and molecular networking.

### Statistical analysis

Statistical analysis was performed on platform MetaboAnalyst (https://www.metaboanalyst.ca/). The feature list obtained through MZmine was reformat and adjusted to the data format required by MetaboAnalyst, then filtered based on QC – filtering features if their RSDs are greater than 25% in QC samples, then normalized with pareto scaling. On that basis, non-supervised multivariate analysis, e.g., principal component analysis (PCA), was performed to understand the general pattern among samples. Furthermore, supervised multivariate data analysis such as partial least squares discriminant analysis (PLS-DA) was performed, and variables of influence on projections (VIP) were obtained. Features with VIP values equal or greater than 1 were considered statistically different substances. In addition, univariate analysis was carried out in a pairwise manner, i.e., p-values were calculated at inter-species level (by comparing samples from *Ophiopogon japonicus* and *Liriope spicata*), and intra-species level (by comparing *L. spicata* from two regions or *O. japonicus* from two regions, respectively).

The list of chemical markers to be identified was generated by integrating VIP and p-values. VIP values greater than 1 and p-values smaller than 0.05 were used.

### Molecular network analysis and computational annotation

A molecular network (MN) was created with the feature-based molecular networking (FBMN) workflow (Nothias et al., 2020) on GNPS (https://gnps.ucsd.edu). The data was filtered by removing all MS/MS fragment ions within +/− 17 Da of the precursor mass. MS/MS spectra were window filtered by choosing only the top 6 fragment ions in the +/− 50 Da window throughout the spectrum. The precursor ion mass and the MS/MS fragment ion tolerances were set to 20 mDa. A MN was then created where edges were filtered to have a cosine score above 0.7 and more than 6 matched peaks. Further, edges between two nodes were kept in the network if and only if each of the nodes appeared in each other respective top 10 most similar nodes. Finally, the maximum size of a molecular family was set to 100, and the lowest scoring edges were removed from molecular families until the molecular family size was below this threshold. The spectra in the MN were then searched against GNPS spectral libraries (Horai et al., 2010; Wang et al., 2016). The library spectra were filtered in the same manner as the input data. All matches kept between network spectra and library spectra were required to have a score above 0.7 and at least 6 matched peaks.

The obtained MN was then subjected to MS2LDA and network annotation propagation (NAP) to performs unsupervised substructure discovery and *in silico* structure annotation (da Silva et al., 2018; Wandy et al., 2018). To enhance chemical structural information within the MN, information from MS2LDA, NAP were incorporated into the network using the GNPS MolNetEnhancer workflow (https://ccms-ucsd.github.io/GNPSDocumentation/molnetenhancer/) on the GNPS website (http://gnps.ucsd.edu) (Ernst et al., 2019). Chemical class annotations were performed using the ClassyFire chemical ontology (Djoumbou Feunang et al., 2016).

Data were visualized in Cytoscape (Version 3.10.0) and R studio.

### Taxonomically informed metabolite annotation

To complement GNPS metabolite annotation, the spectra in the MN were searched against the ISDB-DNP (*in silico* data based-dictionary of natural products) spectral library, and a taxonomically informed scoring approach was applied to improve the annotation process. The results included top five hits of candidate structures (Rutz et al., 2019).

## 3 Results & Discussions

In the present study, to comprehensively investigate the metabolomic profiles of *maidong* produced from different origins, tuberous root samples were collected from four distinct production regions and subjected to untargeted metabolomic profiling. We aimed to obtain an overview of the metabolomic diversity of *maidong* and to identify chemical markers to facilitate quality control. Metabolite variation among *maidong* samples was investigated by multivariate data analysis. MS data were organized as a molecular network (MN) to gain structural information, highlighting structurally related metabolites. The statistical results from the metabolomics were integrated into the MN to identify statistically relevant metabolites. This integrative approach generated informative MN, which we explored to pinpoint potential chemical markers for the quality control of *maidong* plant samples. A schematic paradigm is summarized in Fig. 2.

**Fig. 2.**
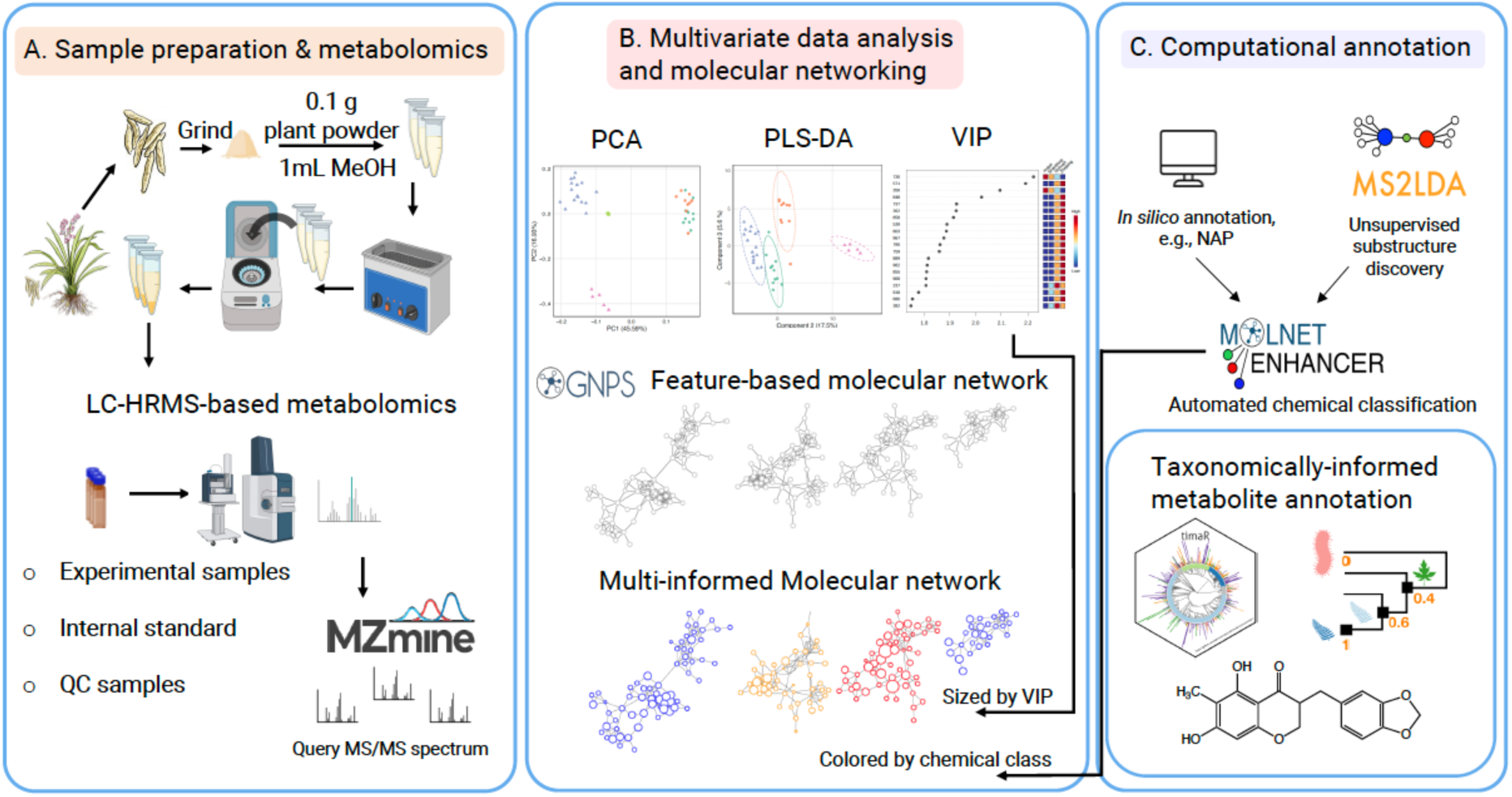
Summary of the schematic paradigm. (A) Sample preparation and LC-HRMS metabolomic analysis. (B) MVDA was applied to investigate the divergences among *maidong*. VIP was used to size the nodes highlighting features of higher importance. (C) Computational approaches were performed to explore chemical classes; metabolite annotation was performed against *in silico* spectral databases, followed by a taxonomic approach.

All extracts prepared from both species were profiled on a single batch by untargeted data-dependent acquisition (DDA) using UHPLC-HRMS/MS and raw data was processed in MZmine for further analysis (Fig. 2A). Initially, a general pattern of metabolomic variation was derived by using unsupervised principal component analysis (PCA). To more precisely identify features statistically important for pattern discrimination, we utilized partial least squares-discriminant analysis (PLS-DA)-a supervised multivariate data analysis (MVDA) model. PLS-DA was selected for its effectiveness in maximizing the correlation between the dataset’s variables and the categorical group assignments, thereby enhancing the discriminative power between different groups. Furthermore, variable importance in projection (VIP) scores was computed within the PLS-DA model to pinpoint compounds with significant discriminatory capability (VIP ≥1). These VIP values are crucial as they prioritize features that contribute most to the model, thus guiding feature selection for further analysis. VIP values and relative concentrations of the compounds were visualized within the MN (Fig. 2B).

In addition, the MS/MS dataset generated by DDA was used as a foundation for building a single feature-based molecular network (FBMN) using the Global Natural Products Social Network platform (GNPS). Metabolite annotation was performed based on comparisons with *in silico* MS/MS fragmentation, and further refined following a taxonomic approach (Fig. 2C).

By integrating MVDA results into the FBMN, we were able to obtain an informative MN facilitating the identification of chemical markers regarding the quality assessment and control over *maidong* derived from different species and regions.

### Metabolomic profiling

In total, 42 *maidong* samples from different regions including *L. spicata* and *O. japonicus* were collected and subjected for metabolomic profiling. A non-targeted metabolomic approach was applied to detect as many features as possible in their metabolome. The data were acquired in both (+)- and (-)-ESI modes utilizing DDA strategy. Extracts were injected in randomized order. In addition, the sequence included QC and blank samples at regular intervals to reduce bias and assess the quality of the metabolomic profiles (Theodoridis et al., 2011).

After the data processing, the dataset yielded 1128 features in (+)-, and 1151 features in (–)-HRMS mode. However, FBMN retrieved on the basis of (–)-ESI data (Fig. S1) provided little valuable information on the presence, for example, of molecules of pharmacological interest such as saponins and flavonoids. This key aspect is intended to facilitate subsequent annotation, which is seen as the essential basis for downstream analysis in this work. Therefore, this study investigated dataset collected in (+)-ESI mode. After processing, the dataset was submitted for MVDA.

Intensities of the internal standard were evaluated to validate the stability and reproducibility of the instrument and data acquisition (Fig. S2), one data point is considered an outlier and thus was removed from further analysis.

### Multivariate data analysis

Unsupervised PCA was carried out to reveal the general pattern of the samples for both species collected in different geographical regions. To ensure the reliability of the data processing, features were filtered out if their RSDs were greater than 25% in QC samples, resulting in a final data set containing 775 features. Pareto scaling and log transformation were applied to reduce the impact of abundant metabolites and address heteroscedasticity by stabilizing the variance and improving the performance of multivariate models.

The PCA model yielded 8 principal components, the first three explaining 68.59% of the variation in the dataset (Fig. 3). This PCA revealed a tight and center clustering of the QC samples, indicating the quality of the dataset is at an acceptable level. Three major groups were assigned, 1) *O. japonicus* produced in Sichuan, 2) *O. japonicus* produced in Zhejiang, 3) *L. spicata*. The distinct chemodiversity between *O. japonicus* produced in the two regions could be associated with different agricultural measures carried out there. *O. japonicus* is cultivated for one year in Sichuan but for three years in Zhejiang (Sun et al., 2020). *L. spicata* produced in two regions were clustered together, suggesting their high similarities in metabolomics. This observation could be credited to their similar agricultural practice, i.e., cultivated for one year in both regions. Besides, the two production regions for *L. spicata* are geographically closer to each other than those for *O. japonicus*. The lower metabolomic divergences resulted in keeping them in the same cluster in the PCA. Therefore, a supervised MVDA, PLS-DA, was performed to focus on metabolites that can better highlight differences between groups, suppressing metabolites that do not contribute significantly to the classification.

**Fig. 3.**
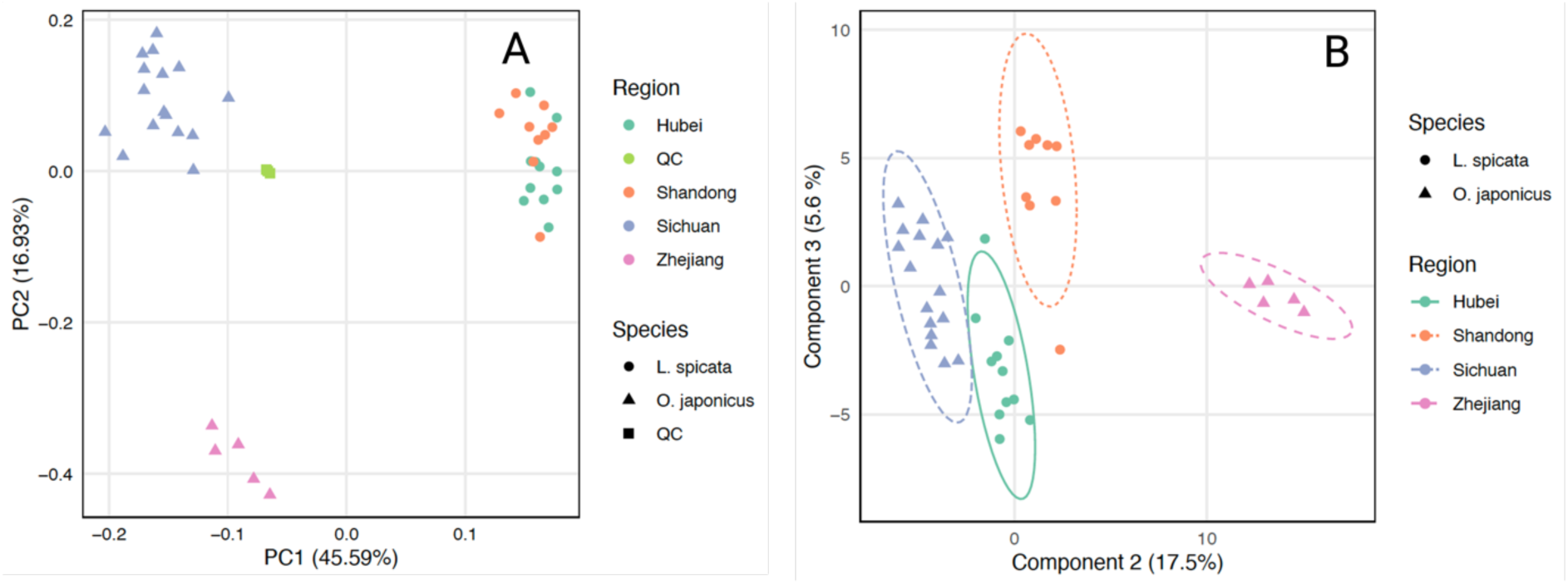
Multivariate data analysis of *maidong* with different botanical and geographical origins (A) PCA score plot of all samples, (B) PLS-DA score plot of the sub-dataset.

### Selection of statistically significant chemical markers through supervised analysis

A supervised PLS-DA was applied to identify the features that were significantly contributing to the difference observed in the PCA plot. Therefore, a sub-dataset without QCs was created for PLS-DA, and an acceptable predictivity was validated by cross-validation showing R^2^= 0.968 and Q^2^= 0.939 for the first three components.

The PLS-DA score plot showed a clear separation among *maidong* from 4 regions on component 2 (Fig. 3B). The chemical markers were selected on the variable importance in the projection (VIP) retrieved from the second component of PLS-DA. VIP value ranged from 0 to 2.2945. Only features with VIP value ≥ 1 were considered statistically significant and thus used for chemical marker selection (Table S3). Eventually, 201 features with VIP values greater than 1 were selected. In the MN, these features were mapped and sized according to their corresponding VIP values, highlighting the features with high discriminating ability.

In addition to the MVDA, a univariate analysis was performed using a t-test to assess the statistical significance of the features at inter- and intra-species levels. In the present study, P-values ≤ 0.05 were considered statistically significant.

### MN analysis and metabolite annotation

The FBMN was generated via the GNPS platform (Nothias et al., 2020). The 1128 features were grouped into 74 clusters based on the similarity of their spectral fragmentation patterns (Fig. 4A, grey MN).

**Fig. 4.**
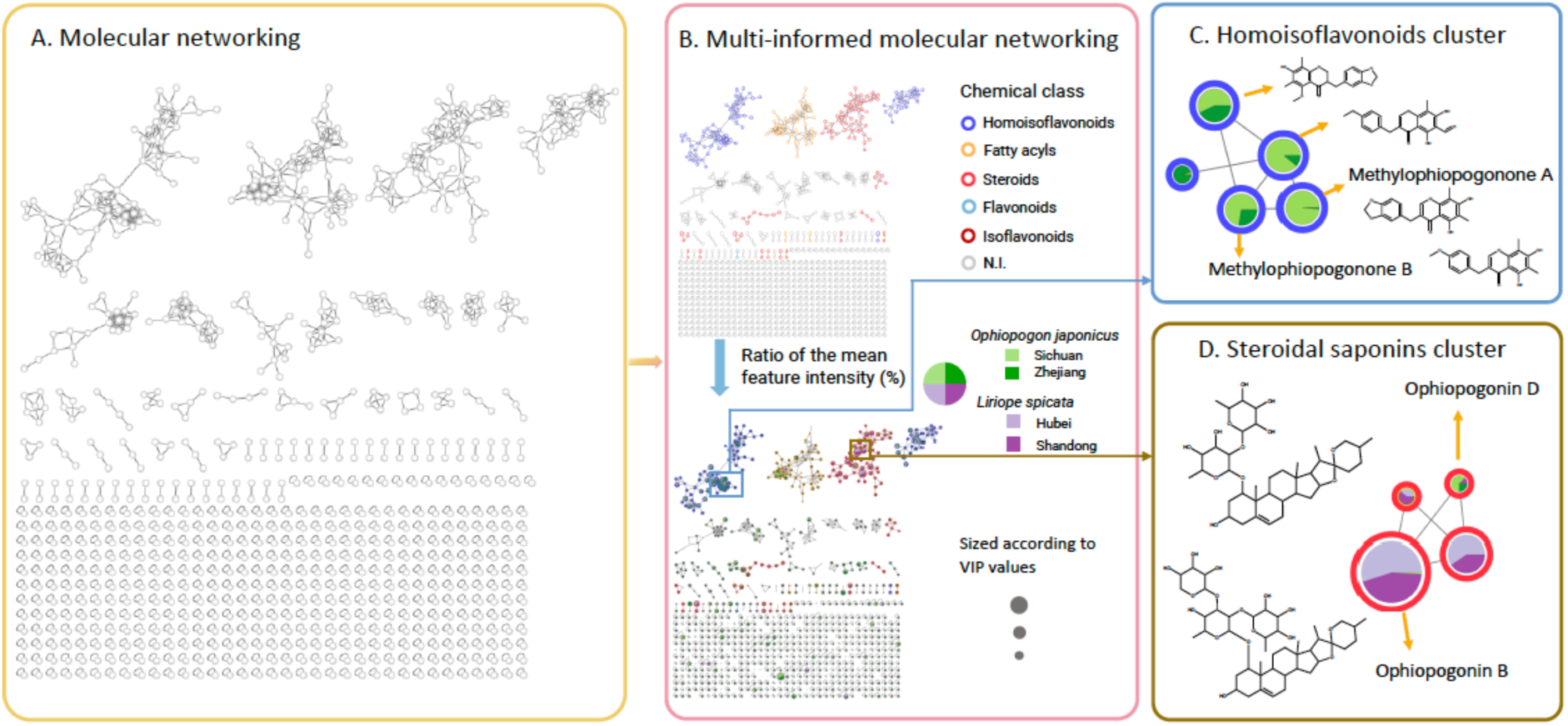
Summary of the procedures used to explore chemical markers (A) MN generated with LC-(+)-HRMS data from *maidong* derived from *O. japonicus* and *L. spicata* in different regions via GNPS. (B) Multi-informed MN featured by chemical classes, and a pie chart inside each node indicating the relative mean intensity of the feature in different groups. FBMN- and MVDA-guided annotation results on homoisoflavonoids (C) and steroidal saponins (D) visualized on MN.

The automated chemical classification of the features was carried out using the MolNetEnhancer algorithm by integrating the results from MS2LDA and NAP analysis and falls in line with ‘ClassyFire’ structure-based ontology (da Silva et al., 2018; Djoumbou Feunang et al., 2016; Wandy et al., 2018). This gave us a holistic view of the primary and secondary metabolites present in *maidong* plants. Furthermore, the MN can be systematically organized and colored at the compound class level based on ClassyFire (Fig. 4B). The main chemical annotated classes were homoisoflavonoids (21.62% out of all features), steroids and steroid derivatives (20.05%), and fatty acyls (14.19 %) (Fig. S3). The direct parent of the steroid features in cluster 2 are steroidal saponins (Fig. S4), we thus refer compounds in cluster 2 as steroidal saponins, and concluded homoisoflavonoids and steroidal saponins are the two dominant chemical classes in *maidong*.

Homoisoflavonoids and steroidal saponins compounds were recognized as characteristic metabolites of *O. japonicus* (Lei et al., 2021). A heatmap (Fig. S5) was generated based on these two classes of compounds. Results indicated that homoisoflavonoids were more pronounced in *O. japonicus* than in *L. spicata* individuals, and *maidong* derived from *O. japonicus*, particularly those from Zhejiang, exhibited a higher level and variety of homoisoflavonoids. In contrast, *maidong* derived from *L. spicata* were enriched in steroidal saponins, with similar metabolomic profiles observed across different production regions.

To better visualize the variation of the relative amounts, the relative mean feature intensities of samples from four regions were displayed as a colored pie chart inside the node of MN (Fig. 4B).

The structural annotation was achieved by running the tima-R script (taxonomically informed metabolomic annotation). In essence, tima-R is used to annotate MS2 spectra by searching against ISDB-DNP (Allard et al., 2016). Subsequently, the annotation output was reranked based on the final score attributed to their biological source, i.e., for which the biological source was found to be “Aspargaceae” at the family level, “*Ophiopogon*” or “*Liriope*” at the genus, and “*Ophiopogon japonicus*” or “*Liriope spicata*” at the species level. Overall, 68.79 % of the features were annotated. Among the 1128 features, 190 (16.84 %) were indicated as having structural candidates found in Asparagaceae family, 91 (8.07%) in ‘*O. japonicus*’ and 8 (0.71%) in *L. spicata*’, respectively (Fig. 5). Fewer features were annotated in *L. spicata* samples, which could be explained by the lack of published data. A total of 47 and 434 hits were found in Web of Science by searching for ‘*Liriope spicata*’ and ‘*Ophiopogon japonicus*’ as keywords, respectively.

**Fig. 5.**
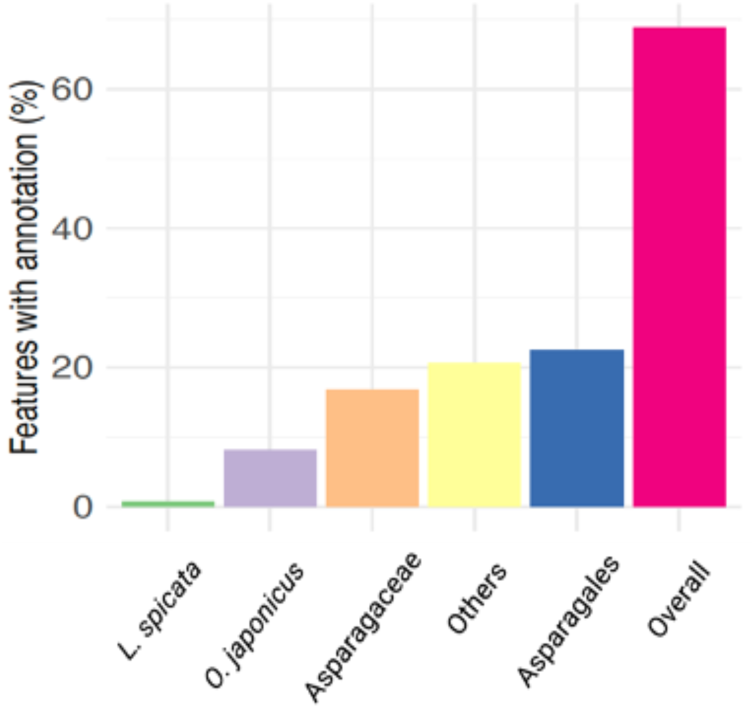
Percentage of features assigned to corresponding taxa computed by the tima-R approach.

### Integration of MVDA with MN

By incorporating statistic data such as VIP and p-values, as well as inserting a pie chart into the MN, we have created a multi-informed MN that provides a comprehensive overview of the chemical classes present in *O. japonicus* and *L. spicata* plants. In addition, clusters of analogues characterizing *O. japonicus* and *L. spicata* were observed at inter-species level, as well as distinctions within each species at the intra-species level. The underlined features could be selected for further detailed analysis (Fig. 4C and D). This integrative approach facilitated a deeper understanding of the chemical diversity of these species, guiding subsequent analysis.

### Selection of clusters and identification of chemical markers

To systematically explore the MN, clusters were selected based on their node size, larger nodes indicating a VIP ≥ 1. Nodes that were not connected to other nodes are called singletons. Singletons were not structurally close to main chemical compound classes, reflecting little valuable information regarding compounds from the main biosynthetic pathway of *O. japonicus* and *L. spicata*. Therefore, they were not further discussed in this study.

After noticing on the MN that homoisoflavonoids and steroidal saponins were the dominant compound classes, the clusters 1-3 were selected for further investigations (Fig. S4). Moreover, compounds from these three groups have been shown to be responsible for a wide range of pharmacological properties of *maidong* (Lei et al., 2021). Structural annotation of features with VIP ≥ 1 and/or p-values ≤ 0.05 were summarized in table S3.

Initially, Cluster 1 and 3 (Fig. S4) were both putatively identified as homoisoflavoinoids (Fig. 6). The key difference between the two clusters was the presence of a double bond in the C2-C3 position (Fig. S6). The majority of metabolites annotated in Cluster 1 contained a double bond at the C2-C3 position corresponding to the class of homoisoflavone derivatives. This double bond was absent in the structures revealed in Cluster 3 and these features were therefore annotated as homoisoflavanones. The majority of the homoisoflavonoids were exclusively found in *O. japonicus*, which is in accordance with the observation that they are characteristic compounds of this plant (Lei et al., 2021).

**Fig. 6.**
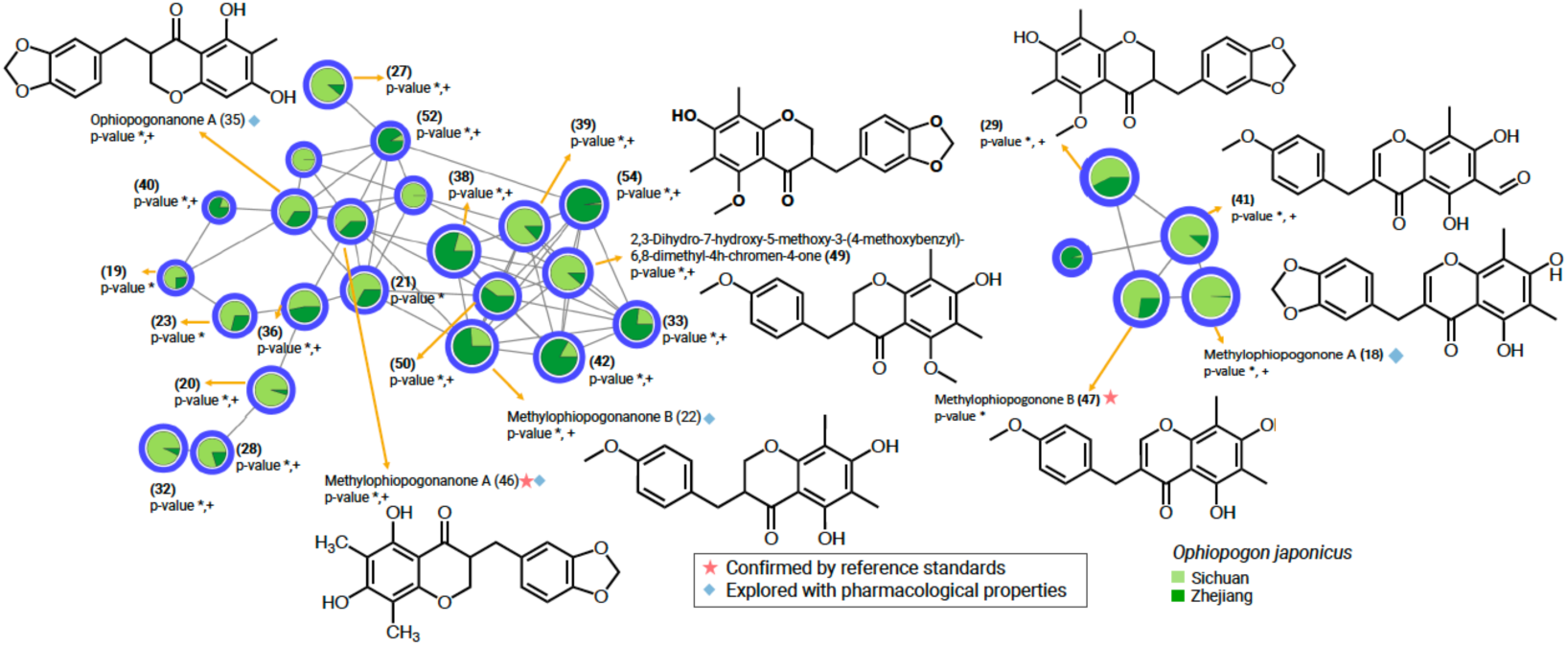
Selected homoisoflavonoid clusters with statistically important features (VIP ≥ 1 and p-value <0.05), i.e. * indicated feature’s p-value < 0.05 at inter-species level. ^+^ indicated feature’s p-value <0.05 at intra-species level for *O. japonicus* ^++^ indicated feature’s p-value <0.05 at intra-species level for *L. spicata*

The MN of some of the steroidal saponins annotated in cluster 2 are shown in Fig. 7. They were detected in both *O. japonicus* and *L. spicata* species. They are most are more abundant in *L. spicata* and only a few are shared by both species. In most cases, each species has its own characteristic steroidal saponins.

**Fig. 7.**
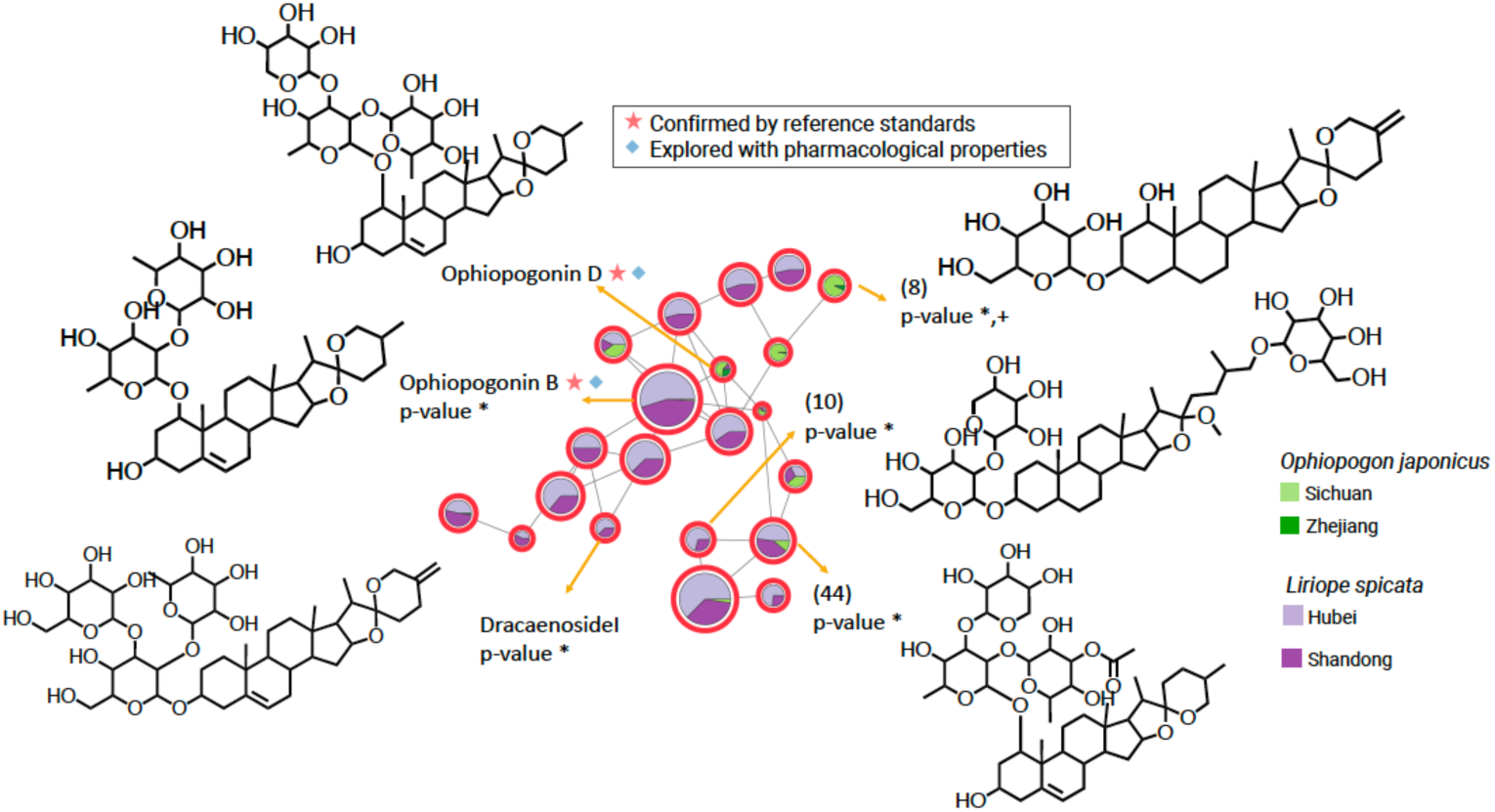
Selected steroidal saponins clusters with high VIP values (p-values were also taken into consideration, i.e. * indicated feature’s p-value < 0.05 at inter-species level + indicated feature’s p-value <0.05 at intra-species level for *O. japonicus* ++ indicated feature’s p-value <0.05 at intra-species level for *L. spicata*

Structures of ophiopogonin D, ophiopogonin B, methylophiopogonanone A, and methylophiopogonone B have been confirmed at level 1 (Schymanski et al., 2014) by comparing the MS2 spectra of putative compounds found in the root extracts with reference standards (MS2 spectra and LC retention times are summarized in Fig. S7). They are often quantified to understand the metabolomic variation of *maidong* derived from different botanical (*O. japonicus* and *L. spicata*) and geographical origins (Wu et al., 2014; Lyu et al., 2020). However, the discussion in these studies is limited to mere quantitative comparison of specific compounds across batches, without highlighting any compounds showed significant variations. In contrast, we employed an untargeted metabolomic approach with several advancements. Our method allowed for the unbiased identification of metabolites without prior selection. Furthermore, MVDA was employed for recognizing molecules with striking contribution to the discrimination of *maidong* derived from different origins. This was achieved by filtering out features with VIP values below 1 and/or p-values above 0.05. Without such meticulous consideration, assessments risk introducing biases and potentially flawed conclusions. Our methodology enhances therefore the reliability of detecting metabolomic differences and contributes to a more accurate understanding of *maidong*’s chemical diversity.

Specifically, in the present study, methylophiopogonanone B regulated by European Pharmacopoeia (Council of Europe, 2023), together with methylophiopogonone B were exclusively found in *O. japonicus*. These two compounds were found to be significantly different between *O. japonicus* cultivated in two regions. This strongly suggested their potential as chemical marker to distinguish *O. japonicus* from *L. spicata* plant extracts, and *O. japonicus* cultivated in two regions. On the other hand, steroidal saponin compounds were found to be ubiquitous as evidenced by their existence across both species from the four regions studied. The levels of ophiopogonin B were observed to be significantly (p-value <0.01) lower in *O. japonicus* compared to *L. spicata,* whereas ophiopogonin D was detected with significantly (p-value < 0.05) higher amounts in *O. japonicus*. Similar results were also found by Chen et al. (2023) in a targeted analysis of *maidong* samples collected in three distinct regions. Different amounts of these two compounds, at intra-species level, were observed in *O. japonicus* and *L. spicata* respectively. However, t-test did not support their contribution at a significantly level, i.e., p-value> 0.05.

Beyond compounds identified at level 1, several compounds were putatively annotated exhibiting significant contribution for pattern recognition. For instance, methylophiopogonone A and ophiopogonanone A have been exclusively found in *O. japonicus*, and could therefore be considered as chemical markers to distinguish *O. japonicus* from *L. spicata* plant extracts. Two homoisoflavonoids derivatives were detected in both species and putatively annotated as 5,7-dihydroxy-3-[(4-hydroxyphenyl)methyl]-2,3-dihydrochromen-4-one and 3-[(3,4-dihydroxyphenyl)methyl]-7-hydroxy-5-methoxy-2,3-dihydrochromen-4-one according to Fig. S4 (in cluster 4). The first compound showed VIP of 1.18, which contribute to differentiate in the PLS-DA model between *maidong* harvested in the four regions. Meanwhile, the t-test confirmed the significant ability of discriminating these two homoisoflavonoids at inter-species, but not at intra-species levels. The second derivative (VIP< 1) was excluded when running PLS-DA processing due to its low power to discriminate *maidong* harvested in different regions.

To summarize, at inter-species level, *O. japonicus* and *L. spicata* differed in their steroidal saponins as well as flavonoids composition. Homoisoflavonoids were shown to be characteristic metabolites in *O. japonicus* but lack in *L. spicata*, steroidal saponins presented more in *L. spicata* rather than *O. japonicus*. At the intra-species level, *O. japonicus* collected in two regions tended to have two chemotypes and several metabolites were putatively identified as chemical markers to differentiate them. This could be attributed to their different growing times and geographical distribution (Ge et al., 2019). However, *L. spicata* collected in two regions were very close to each other, and only a few metabolites were considered to be significantly different.

### Analysis of commercial materials

From a commercial point of view, *O. japonicus* from Zhejiang is the subject of exclusive supply agreements under which it is sold directly to specific companies, thus bypassing the traditional medicinal market. Meanwhile, *L. spicata* produced in Hubei and Shandong is often grouped together under the general label of *L. spicata* products, without the emphasis being placed on the region of production. Moreover, (in)advert mix of different origins samples can happen during the trade of medicinal herbs (Zhang et al., 2012). These practices result in a significant heterogenization of the metabolite profiles, hampering quality control efforts. Consequently, commercially traded *maidong* samples were also investigated to determine if our analytical approach could be adapted to these more complex scenarios.

Commercial samples of *maidong* from *O. japonicus* and *L. spicata* roots have been profiled, and the metabolomic data was analyzed following the afore-introduced procedure (Fig. 2). MVDA was performed to obtain an overview of their metabolomic profile. The first two PC levels of the resulting PCA explained 46.55% of the variance (Fig. 8). The metabolite composition of *O. japonicus* and *L. spicata* samples were found to be significantly different at the inter-species level and can be distinguished by unsupervised model PCA. Three suspicious samples declared to be *O. japonicus* were grouped with *L. spicata*, and could therefore be showcases of adulteration. Meanwhile, one sample declaring *O. japonicus* as the botanical source was statistically considered as an outlier. Thus, the four potential mislabeling *maidong* samples and the outlier were removed when calculated mean intensity of the features.

**Fig. 8.**
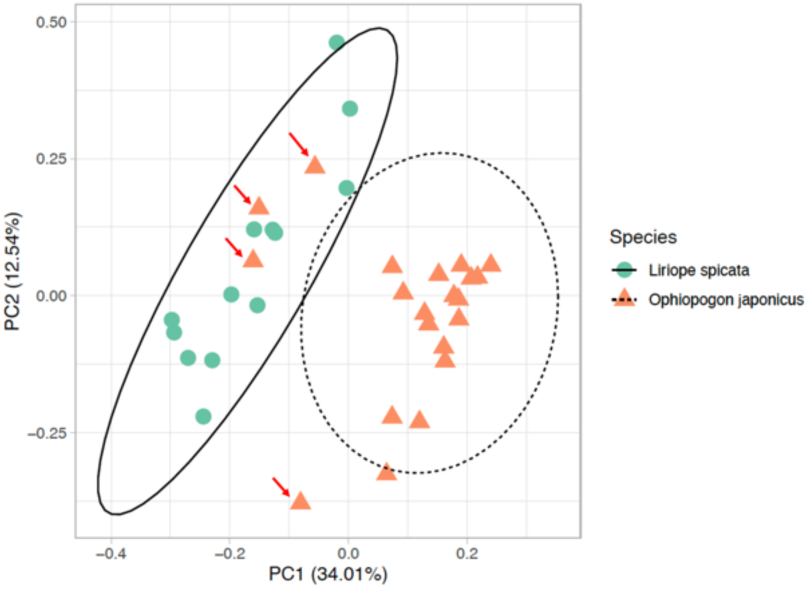
PCA score plot of *maidong* samples purchased in markets in China.

Specific differences in the composition of steroidal saponins and homoisoflavonoids in *maidong* from *O. japonicus* and *L. spicata* were also examined. A MN was generated with 410 nodes divided in 28 clusters (Fig. S8). The results are in line with our observations on *maidong* collected from fields, i.e., the exclusive existence of homoisoflavonoids in *O. japonicus* was confirmed, as well as higher levels of Ophiopogonin D in *O. japonicus* (Fig. 9). This means, chemical markers such as ophiopogonin D, ophiopogonin B, methylophiopogonone B, and methylophiopogonanone B are also very different in commercial samples.

**Fig. 9.**
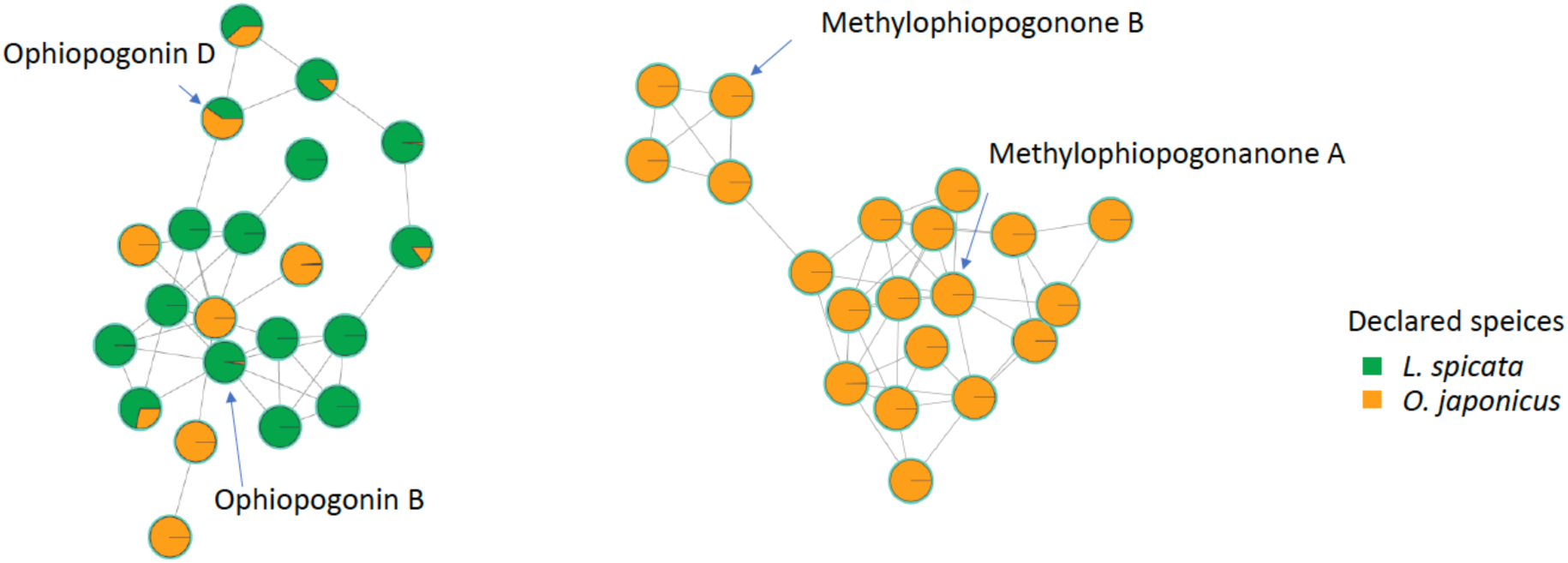
Selected cluster of steroidal saponins and homoisoflavonoids in the MN of commercial *maidong* root samples.

In summary, this untargeted metabolomic study showcased a holistic overview on the metabolomic diversity and differences among *maidong* derived from different origins. The MN demonstrated that steroidal saponins and homoisoflavonoids to be the dominant chemical classes in *maidong*. Our results showcased the distinct metabolites among *maidong* derived from four origins, i.e., exclusive presence of broad ranges of homoisoflavonoids in *O. japonicus*, whereas more exclusive presence of steroidal saponins in *L. spicata*. Overall, 58 metabolites that substantially contributed to the inter-specific discrimination were annotated. It emerged that 6 and 36 metabolites respectively were annotated as being significant in discriminating between *L. spicata* and *O. japonicus* at a regional level. This will enable further studies to be carry out to investigate their pharmacological properties in a more targeted way. These compounds could potentially facilitate the standardization of *maidong* quality control and contribute to more sustainable production and use, as well as more rigorous quality control.

Finally, a wide range of primary metabolites such as fatty acyls were detected in the present study. Unfortunately, little pharmacological research has investigated on this class of chemical components. Future studies may take fatty acyls into consideration when evaluating their therapeutic efficacy for a more comprehensive understanding of medicinal plants such as *maidong* (Hou et al., 2019).

## 4 Conclusion

A comprehensible view on the metabolomic diversity and variation of *maidong* was achieved through the untargeted metabolomic analysis using LC-HRMS. Data interpretation was facilitated by integration of standard uni- and multivariate data analysis and molecular networking. This approach enabled clear differentiation of *maidong* from different origins, highlighting the contributing inter- and intra-specific features. With the support of *in silico* annotation tools, the chemical structure of some of the features could be classified and even elucidated. Steroidal saponins and homoisoflavonoids were recognized as the predominant classes of compounds through employing a more systematic approach, rather than focusing on a specific subset of metabolites.

Additionally, our methodology has also proved to be suitable for quality control of commercial *maidong* samples. However, our understanding on the metabolites present in *L. spicata* is constrained by the scarcity of MS data of compounds of this species and accessible authentic references. This coud be improved with more extraction, purification, and structure elucidation work targeting this species.

The present study offers insights into selecting chemical markers for quality assessment by fostering a holistic understanding on metabolomic diversity and variation of the organisms. This approach is beneficial for metabolomic studies where *a prior* knowledge on the metabolomic profiles of the organism is limited, and a hypothesis-generating experiment is required. This is particularly important when plant-based medicines derived from multiple botanical origins. Consequently, this approach ensures a more rigorous selection of pharmacologically relevant natural ingredients that can be used in phytomedicine.

## Supporting information

maidong_FBMN_supplment

## Data Availability

FBMN generated can be downloaded from https://gnps.ucsd.edu/ProteoSAFe/status.jsp?task=c57030cae2044a5d831ef712529de955 and https://gnps.ucsd.edu/ProteoSAFe/status.jsp?task=1c71501cddb246dc92870f9d6fadca0e. MolNetEnhancer results can be downloaded from https://gnps.ucsd.edu/ProteoSAFe/status.jsp?task=463c77e042634b02bd2001167a09abe8.

## CRediT authorship contribution statement

Lei Feiyi: Conceptualization, Data curation, Investigation, Methodology, Visualization, Writing – original draft, Writing – review & editing. Saldanha L.Leonardo: Data curation, Writing-review & editing. Weckerle Caroline Sonja: Conceptualization, Writing – review & editing. Bigler Laurent: Metabolomics, writing-review & editing.

## Acknowledgement

We thank Dr. Reto Nyffeler (Zurich) for his contribution on botanical identification, Prof. Xingfu Chen, Dr. Tao Wang, Ms. Cuirong Zhao, Mr. Kezhao Liu, Mr. Ticai Wang, Mr. Jianzhong Xu, Ms. Sijia Li for their generous help on sample collection in the field.

