## Supplementary material for "Discovery of chemical marker for *maidong* (roots of *Ophiopogon japonicus* and *Liriope spicata*): a feature-based molecular networking approach": maidong_FBMN_supplment

**Fig. S1** FBMN of *maidong* obtained from LC-MS data acquired in the negative ionization mode.

**Fig.** **S2** Evaluation of the stability of the LC-MS instrument and the reproducibility of data acquisition.

**Fig.** **S3** Pie chart of features sorted according to their chemical class assignment and classified by the MolNetEnhancer algorithm.
**Fig.** **S4** Overview of the MN generated based on metabolomic data of *maidong* derived from *O. japonicus* and *L. spicata* harvested in different regions.

**Fig. S5** Heatmap of features annotated as homoisoflavonoids and steroidal saponins.

**Fig. S6** Illustrations of two types of homoisoflavonoids- homoisoflavones and homoisoflavanones found in *maidong*.

**Fig.** **S7** Structure elucidation of ophiopogonin B and D, methylophiopogonone B and methylophiopogonanone A by comparing the MS2 spectra from *maidong* extracts with those of reference standards

**Fig. S8** Overview of the MN generated based on metabolomic data of *maidong* purchased in markets.

**Table S1** Sample list of *maidong* collected in the production regions in China.

**Table S2** Sample list of commercial *maidong* collected in medicinal markets in China.

**Table S3** Identification of compounds in *maidong* samples by UHPLC-HRMS analysis highlighted as inter- and intra-species characteristic metabolites metabolites.

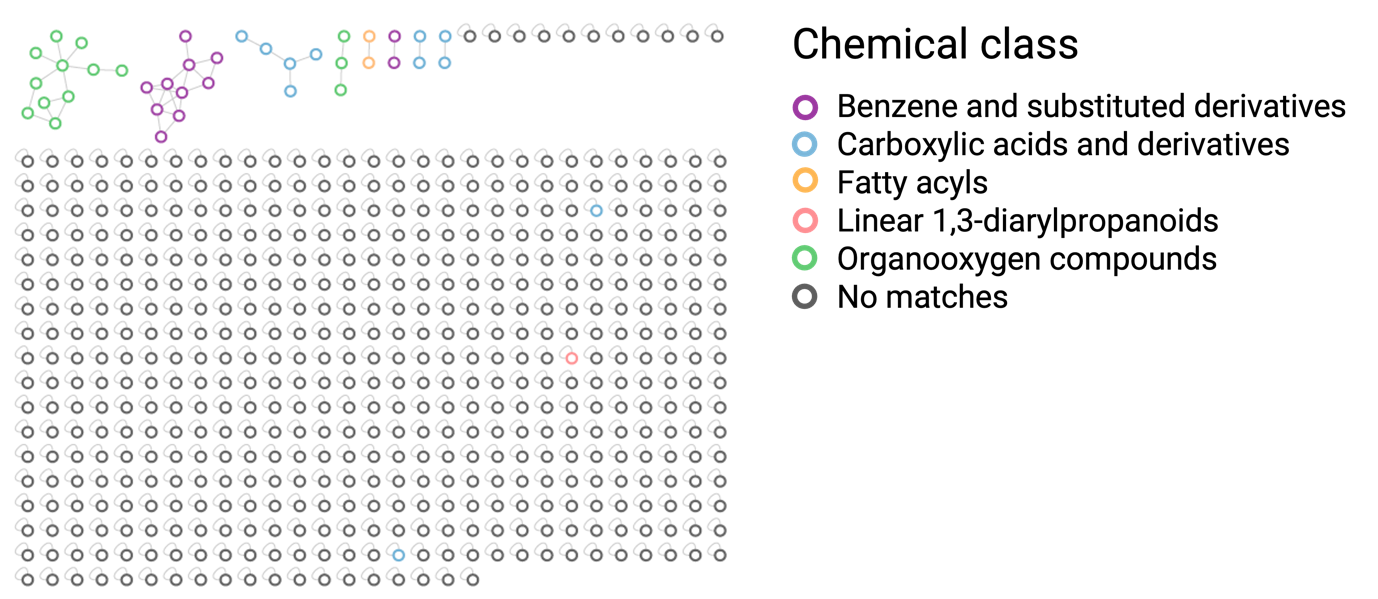

**Fig. S1** FBMN of *maidong* obtained from LC-MS data acquired in the negative ionization mode.

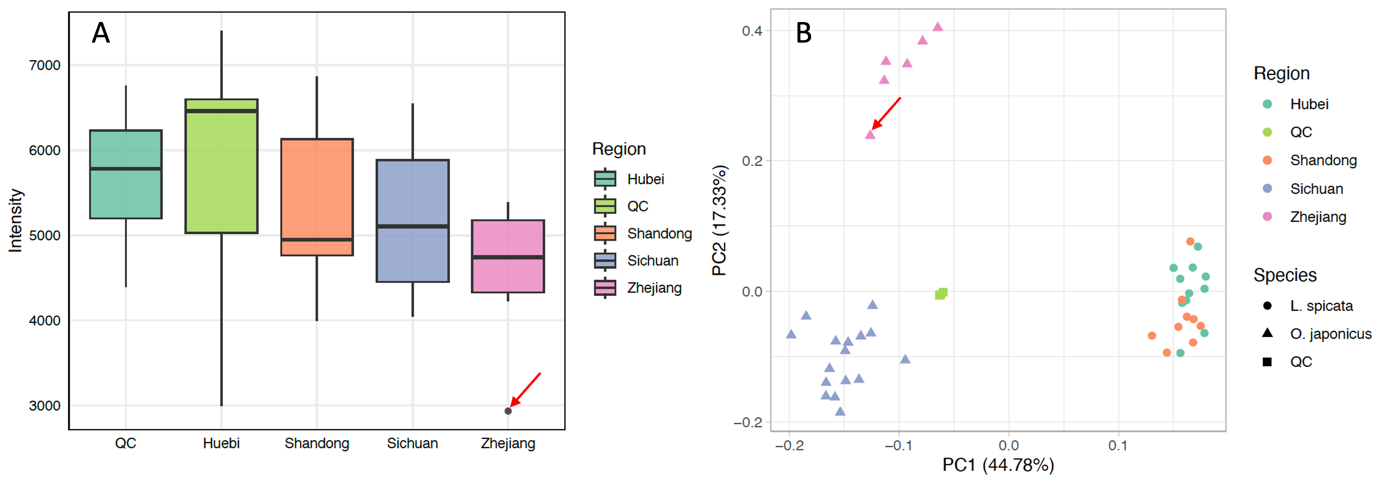

**Fig. S2** Evaluation of stability of the LC-MS instrumentation and the reproducibility of data acquisition. (A) Boxplot of intensity of internal standard, Ampicillin (*m/z* 349.1096 ) in different types of samples; (B) PCA score plot of *maidong* produced in four regions and QC samples. The statistical outlier is indicated by a red arrowed in both plots.

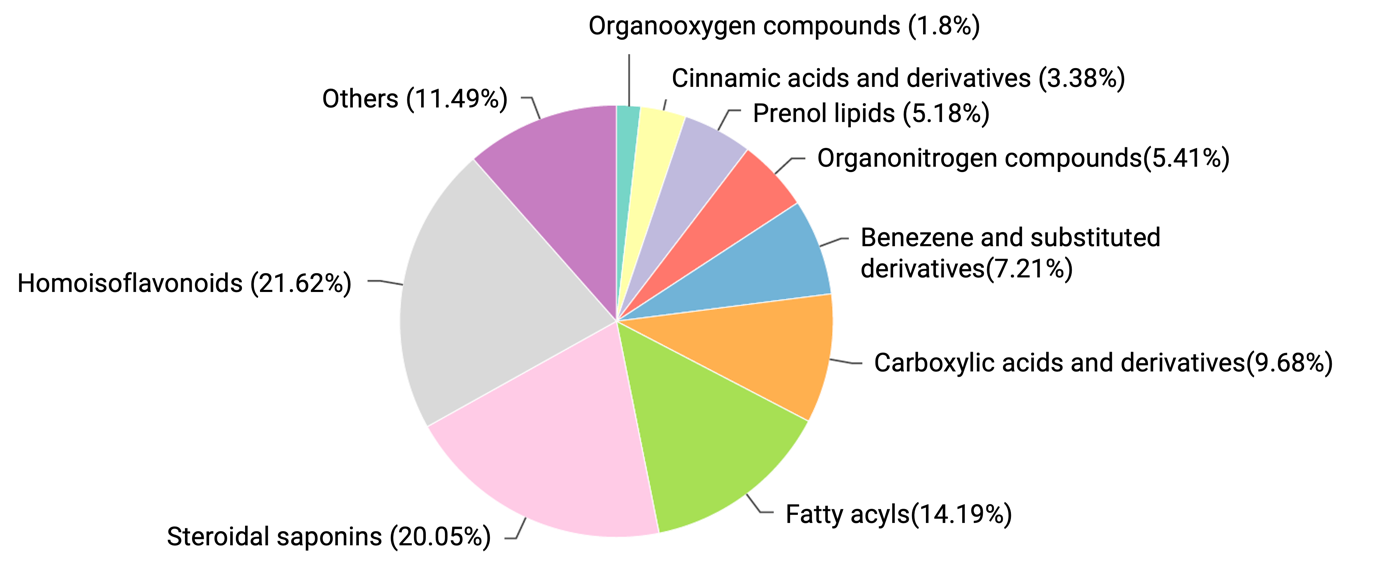

**Fig. S3** Pie chart of features sorted according to their chemical class assignment and classified by the MolNetEnhancer algorithm

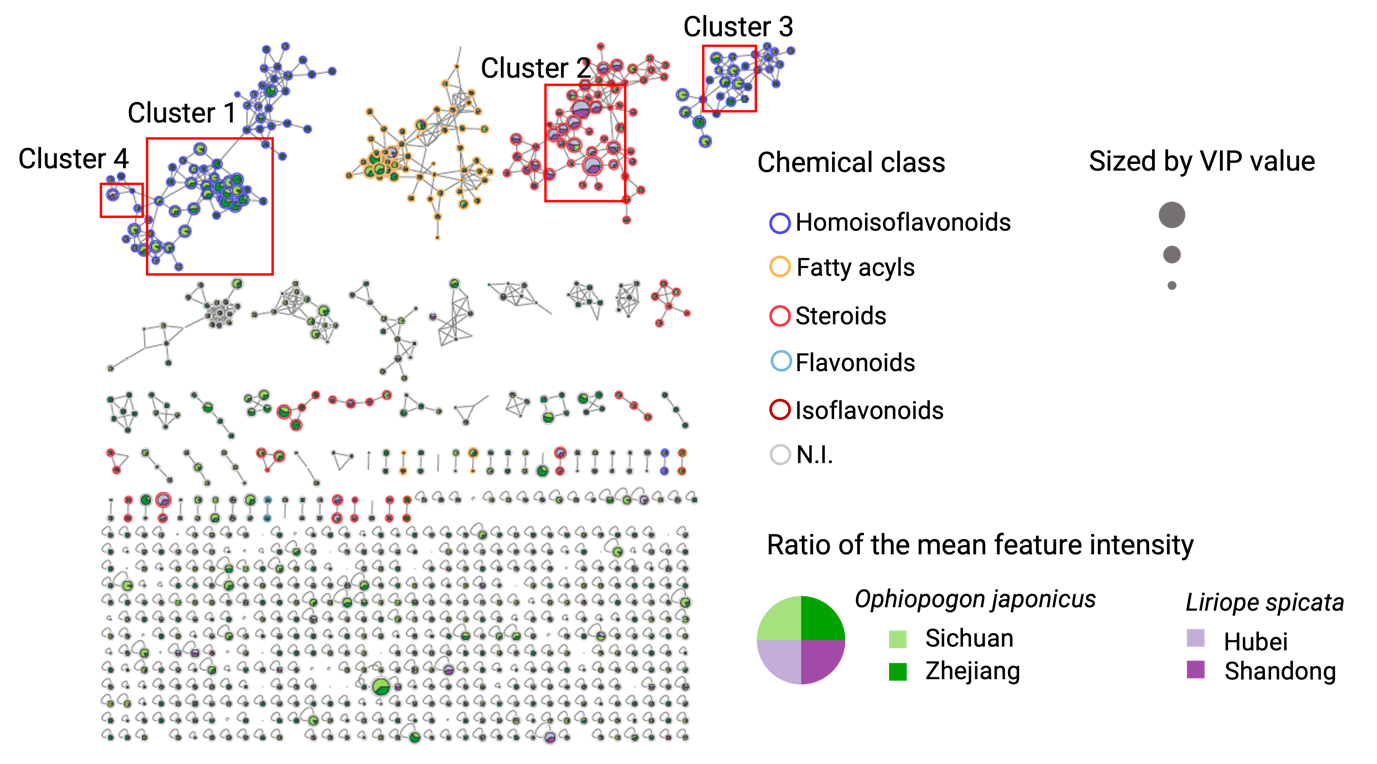

**Fig. S4** Overview of the MN generated based on metabolomic data of *maidong* derived from *O. japonicus* and *L. spicata* harvested in different regions.

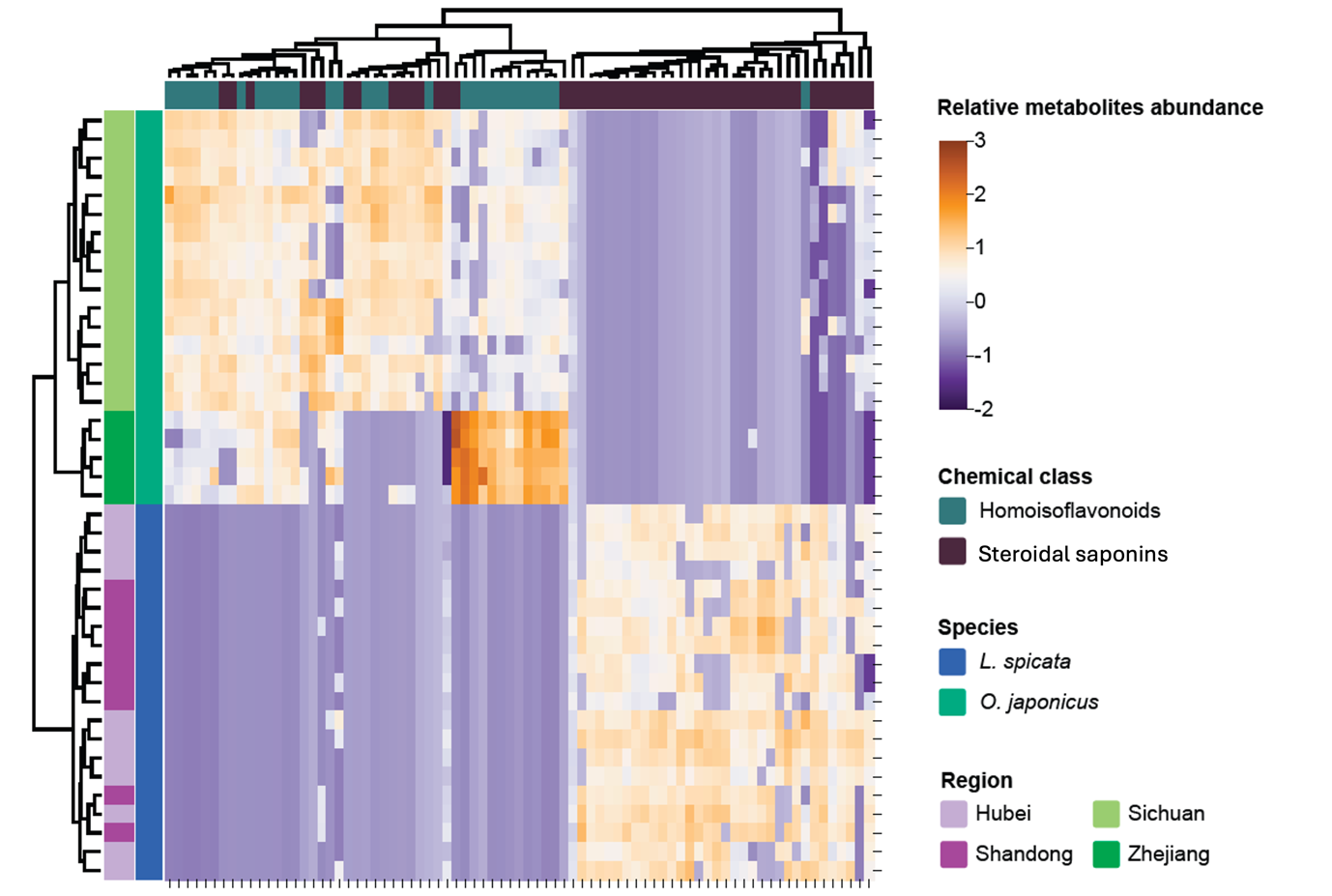

**Fig. S5** Heatmap of features annotated as homoisoflavonoids and steroidal saponins (clustered based on chemical classes, botanical species and harvest regions)

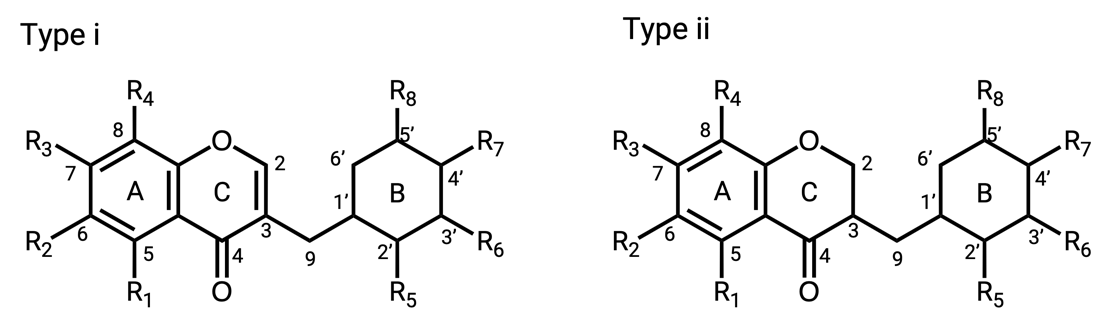

**Fig. S6** Illustrations of two types of homoisoflavonoids- homoisoflavones and homoisoflavanones found in *maidong*.

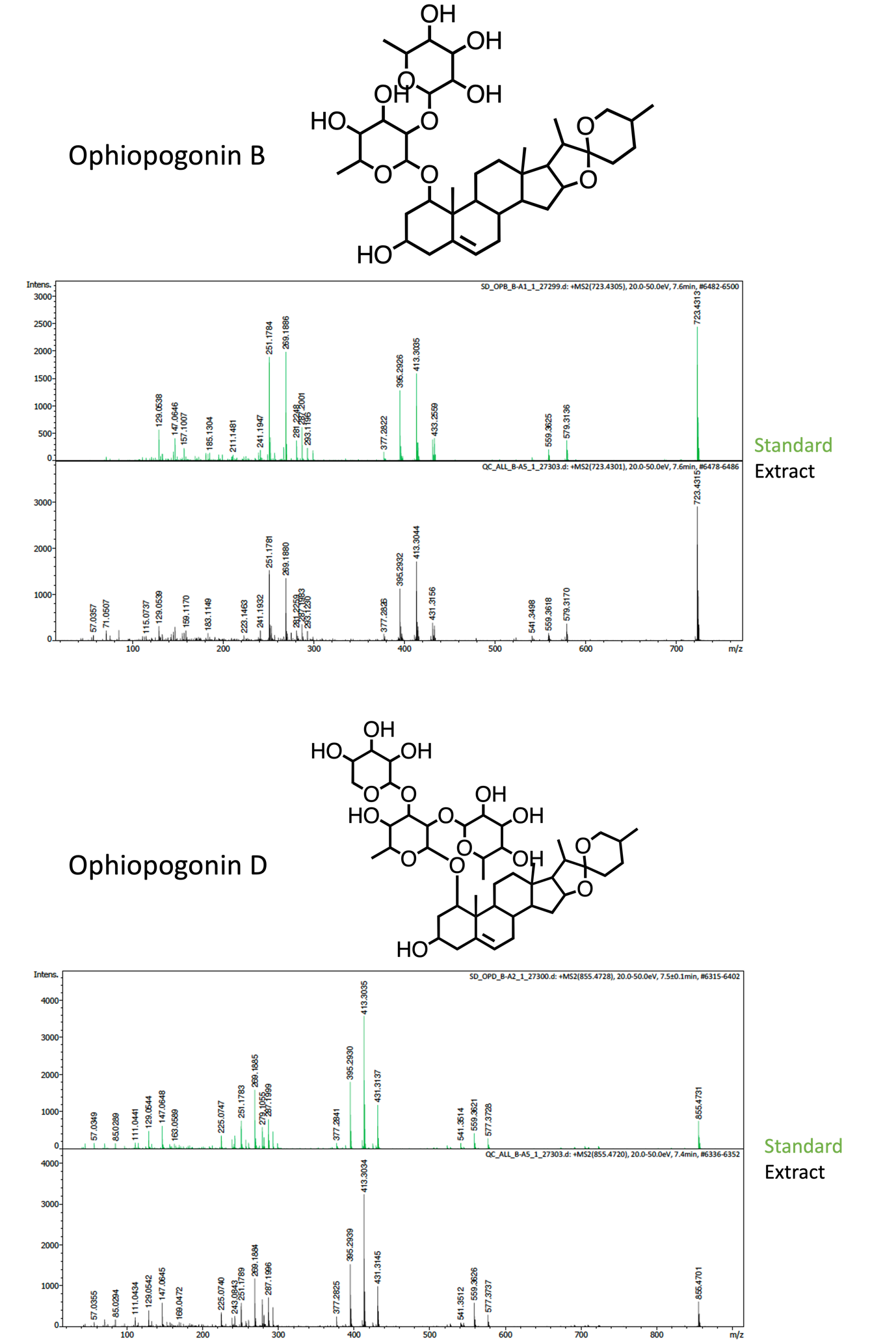

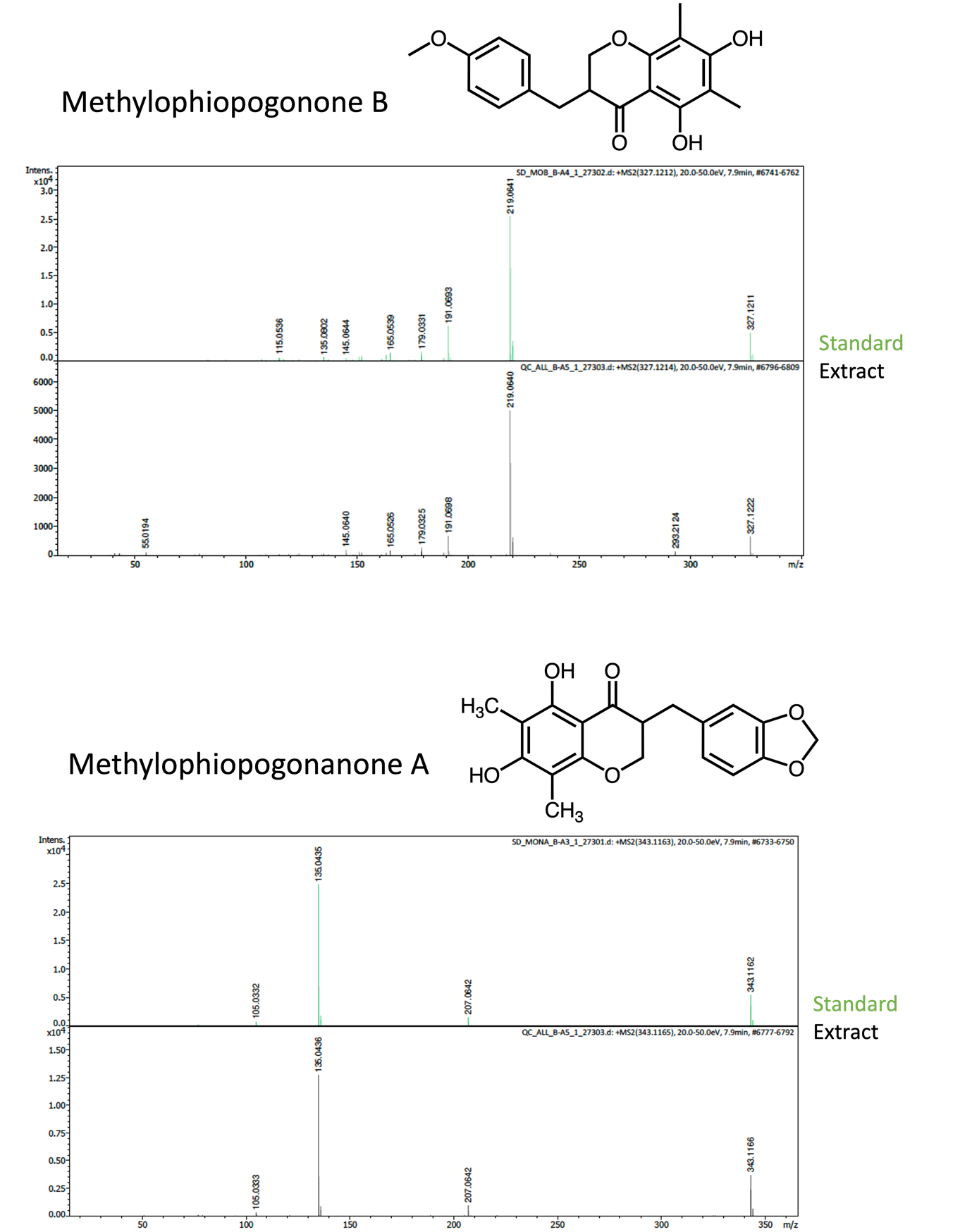

**Fig. S7** Structure elucidation of ophiopogonin B and D, methylophiopogonone B and methylophiopogonanone A by comparing the MS2 spectra from *maidong* extracts with those of reference standards

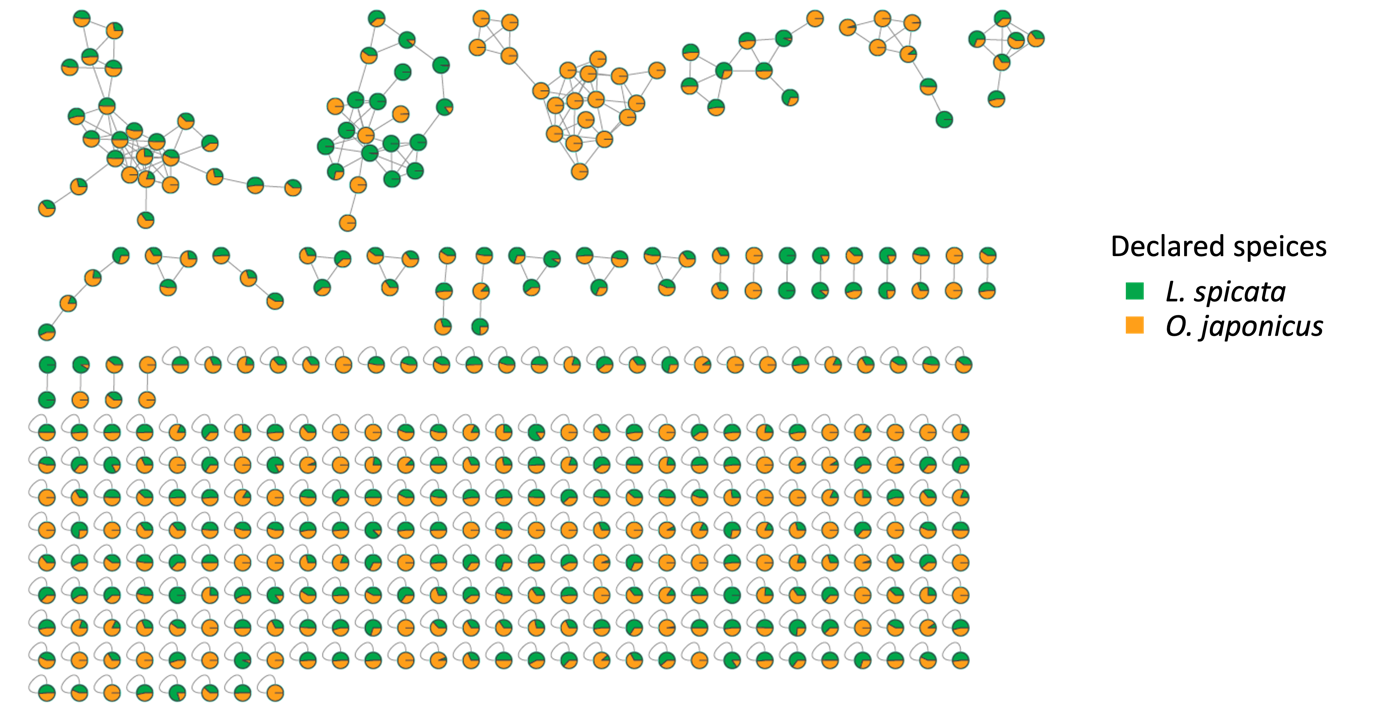

**Fig. S8** Overview of the MN generated based on metabolomic data of *maidong* purchased in markets.

**Table S1** Sample list of *maidong* collected in the production regions in China

| NO. | Species | Region (province) | Root type |
| --- | --- | --- | --- |
| 1 | *Liriope spicata* | Hubei | Tuberous roots |
| 2 | *Liriope spicata* | Hubei | Tuberous roots |
| 3 | *Liriope spicata* | Hubei | Tuberous roots |
| 4 | *Liriope spicata* | Hubei | Tuberous roots |
| 5 | *Liriope spicata* | Hubei | Tuberous roots |
| 6 | *Liriope spicata* | Hubei | Tuberous roots |
| 7 | *Liriope spicata* | Hubei | Tuberous roots |
| 8 | *Liriope spicata* | Hubei | Tuberous roots |
| 9 | *Liriope spicata* | Hubei | Tuberous roots |
| 10 | *Liriope spicata* | Hubei | Tuberous roots |
| 11 | *Liriope spicata* | Hubei | Tuberous roots |
| 12 | *Liriope spicata* | Shandong | Tuberous roots |
| 13 | *Liriope spicata* | Shandong | Tuberous roots |
| 14 | *Liriope spicata* | Shandong | Tuberous roots |
| 15 | *Liriope spicata* | Shandong | Tuberous roots |
| 16 | *Liriope spicata* | Shandong | Tuberous roots |
| 17 | *Liriope spicata* | Shandong | Tuberous roots |
| 18 | *Liriope spicata* | Shandong | Tuberous roots |
| 19 | *Liriope spicata* | Shandong | Tuberous roots |
| 20 | *Liriope spicata* | Shandong | Tuberous roots |
| 21 | *Liriope spicata* | Shandong | Tuberous roots |
| 22 | *Ophiopogon japonicus* | Sichuan | Tuberous roots |
| 23 | *Ophiopogon japonicus* | Sichuan | Tuberous roots |
| 24 | *Ophiopogon japonicus* | Sichuan | Tuberous roots |
| 25 | *Ophiopogon japonicus* | Sichuan | Tuberous roots |
| 26 | *Ophiopogon japonicus* | Sichuan | Tuberous roots |
| 27 | *Ophiopogon japonicus* | Sichuan | Tuberous roots |
| 28 | *Ophiopogon japonicus* | Sichuan | Tuberous roots |
| 29 | *Ophiopogon japonicus* | Sichuan | Tuberous roots |
| 30 | *Ophiopogon japonicus* | Sichuan | Tuberous roots |
| 31 | *Ophiopogon japonicus* | Sichuan | Tuberous roots |
| 32 | *Ophiopogon japonicus* | Sichuan | Tuberous roots |
| 33 | *Ophiopogon japonicus* | Sichuan | Tuberous roots |
| 34 | *Ophiopogon japonicus* | Sichuan | Tuberous roots |
| 35 | *Ophiopogon japonicus* | Sichuan | Tuberous roots |
| 36 | *Ophiopogon japonicus* | Sichuan | Tuberous roots |
| 37 | *Ophiopogon japonicus* | Zhejiang | Tuberous roots |
| 38 | *Ophiopogon japonicus* | Zhejiang | Tuberous roots |
| 39 | *Ophiopogon japonicus* | Zhejiang | Tuberous roots |
| 40 | *Ophiopogon japonicus* | Zhejiang | Tuberous roots |
| 41 | *Ophiopogon japonicus* | Zhejiang | Tuberous roots |
| 42 | *Ophiopogon japonicus* | Zhejiang | Tuberous roots |

**Table S2** Sample list of commercial *maidong* collected in medicinal markets in China

| No. | Declared Species | Root type | Market type | Location |
| --- | --- | --- | --- | --- |
| 1 | *Ophiopogon japonicus* | Tuberous roots | Local medicinal market | Chengdu, China |
| 2 | *Ophiopogon japonicus* | Tuberous roots | Local medicinal market | Chongqing, China |
| 3 | *Ophiopogon japonicus* | Tuberous roots | National medicinal market | Bozhou, China |
| 4 | *Liriope spicata* | Tuberous roots | National medicinal market | Bozhou, China |
| 5 | *Ophiopogon japonicus* | Tuberous roots | National medicinal market | Bozhou, China |
| 6 | *Liriope spicata* | Tuberous roots | National medicinal market | Bozhou, China |
| 7 | *Ophiopogon japonicus* | Tuberous roots | National medicinal market | Anguo, China |
| 8 | *Ophiopogon japonicus* | Tuberous roots | National medicinal market | Anguo, China |
| 9 | *Ophiopogon japonicus* | Tuberous roots | National medicinal market | Anguo, China |
| 10 | *Ophiopogon japonicus* | Tuberous roots | National medicinal market | Anguo, China |
| 11 | *Ophiopogon japonicus* | Tuberous roots | National medicinal market | Anguo, China |
| 12 | *Ophiopogon japonicus* | Tuberous roots | National medicinal market | Anguo, China |
| 13 | *Liriope spicata* | Tuberous roots | National medicinal market | Anguo, China |
| 14 | *Liriope spicata* | Tuberous roots | National medicinal market | Anguo, China |
| 15 | *Liriope spicata* | Tuberous roots | National medicinal market | Yuzhou, China |
| 16 | *Ophiopogon japonicus* | Tuberous roots | National medicinal market | Yuzhou, China |
| 17 | *Liriope spicata* | Tuberous roots | National medicinal market | Yuzhou, China |
| 18 | *Ophiopogon japonicus* | Tuberous roots | National medicinal market | Yuzhou, China |
| 19 | *Liriope spicata* | Tuberous roots | National medicinal market | Yuzhou, China |
| 20 | *Liriope spicata* | Tuberous roots | National medicinal market | Yuzhou, China |
| 21 | *Liriope spicata* | Tuberous roots | National medicinal market | Yuzhou, China |
| 22 | *Ophiopogon japonicus* | Tuberous roots | National medicinal market | Chengdu, China |
| 23 | *Ophiopogon japonicus* | Tuberous roots | National medicinal market | Chengdu, China |
| 24 | *Ophiopogon japonicus* | Tuberous roots | National medicinal market | Chengdu, China |
| 25 | *Ophiopogon japonicus* | Tuberous roots | National medicinal market | Chengdu, China |
| 26 | *Ophiopogon japonicus* | Tuberous roots | National medicinal market | Chengdu, China |
| 27 | *Ophiopogon japonicus* | Tuberous roots | National medicinal market | Chengdu, China |
| 28 | *Liriope spicata* | Tuberous roots | National medicinal market | Chengdu, China |
| 29 | *Ophiopogon japonicus* | Tuberous roots | National medicinal market | Chengdu, China |
| 30 | *Ophiopogon japonicus* | Tuberous roots | National medicinal market | Chengdu, China |
| 31 | *Liriope spicata* | Tuberous roots | National medicinal market | Chengdu, China |
| 32 | *Ophiopogon japonicus* | Tuberous roots | National medicinal market | Chengdu, China |
| 33 | *Liriope spicata* | Tuberous roots | National medicinal market | Chengdu, China |
| 34 | *Liriope spicata* | Tuberous roots | National medicinal market | Chengdu, China |

**Table S3** Identification of compounds in *maidong* samples by UHPLC-HRMS analysis highlighted as inter- and intra-species characteristic metabolites metabolites. Compounds were annotated by means of MS spectra comparison against the ISDB-DNP spectra database following a taxonomic approach. Authentic standards were used as reference compounds to confirm the annotated compounds when available. The partial InChIkey (International Chemical Identifier for the annotated metabolites are presented. VIP values from the PLS-DAare presented. P-values at inter- and intra-species level are presented.

| No. | VIP | Feature ID | Chemical class | Annotation (Correspond compounds in NPD) | pValue_  interspecies | pValue_  IntraOphiopogon | pValue_  IntraLiriope | InChIKey |
| --- | --- | --- | --- | --- | --- | --- | --- | --- |
| 1 | 1.14 | 31 | Steroids and steroidal derivative | Guanidine | 0.009504932 | 5.77E-08 | >0.05 | ZRALSGWEFCBTJO |
| 2 | 1.56 | 310 | homoisoflavonoids | Chromone | 4.79E-14 | 1.25E-05 | >0.05 | OTAFHZMPRISVEM |
| 3 | 1.24 | 329 | Steroids and steroidal derivative | (1R,2S,4S,5'R,6R,7S,8R,9S,12S,13S,16S,18S)- 16-[(2R,3R,4R,5R,6R)-3,4-dihydroxy-5-[(2S,3R,4S,5R,6R)-5-hydroxy-6-(hydroxymethyl)-3,4-bis[[(2S,3R,4S,5S,6R)-3,4,5-trihydroxy-6-(hydroxymethyl)oxan-2-yl]oxy]oxan-2-yl]oxy-6-(hydroxymethyl)oxan-2-yl]oxy-5',7,9,13-tetramethylspiro[5-oxapentacyclo[10.8.0.02,9.04,8.013,18]icosane-6,2'-oxane]-10-one | 3.38E-10 | 1.70E-08 | >0.05 | SQQNDULGFFHXND |
| 4 | 1.16 | 332 | Steroids and steroidal derivative | Filiasparoside A | 1.31E-06 | >0.05 | 2.03E-06 | OUCDJSWSAODKNK |
| 5 | 1.37 | 335 | Steroids and steroidal derivative | 16beta-Acetoxyproscillaridin A | 4.33E-14 | >0.05 | >0.05 | VMLMEZINUVEFME |
| 6 | 1.35 | 336 | Steroids and steroidal derivative | (2S,3R,4R,5R,6S)-2-[(2R,3S,4S,5R,6R)-4- hydroxy-2-(hydroxymethyl)-5-[(2S,3R,4S,5S,6R)-3,4,5-trihydroxy-6-(hydroxymethyl)oxan-2-yl]oxy-6-[(1R,2S,4S,6R,7S,8R,9S,12S,13S,16S,18R)-7,9,13-trimethyl-5'-methylidenespiro[5-oxapentacyclo[10.8.0.02,9.04,8.013,18]icosane-6,2'-oxane]-16-yl]oxyoxan-3-yl]oxy-6-methyloxane-3,4,5-triol | 3.83E-16 | >0.05 | >0.05 | GTHFQEMYVCDXKR |
| 7 | 1.54 | 337 | Steroids and steroidal derivative | Chonglou Saponin VII | 3.53E-15 | >0.05 | 0.00460277 | FBFJAXUYHGSVFN |
| 8 | 1.25 | 345 | Steroids and steroidal derivative | (2S,3R,4S,5S,6R)-2-[(2S,3R,4S,5S,6R)- 2-[(2R,3S,4R,5R,6R)-4,5-dihydroxy-2-(hydroxymethyl)-6-[(1S,2S,4S,5'R,6R,7S,8R,9S,12S,13S,14R,16R,18S)-14-hydroxy-5',7,9,13-tetramethylspiro[5-oxapentacyclo[10.8.0.02,9.04,8.013,18]icosane-6,2'-oxane]-16-yl]oxyoxan-3-yl]oxy-4,5-dihydroxy-6-(hydroxymethyl)oxan-3-yl]oxy-6-(hydroxymethyl)oxane-3,4,5-triol | 4.56E-10 | 1.11E-08 | >0.05 | BPCZJBXLAYHPQL |
| 9 | 1.184 | 348 | Steroids and steroidal derivative | DracaenosideI | 7.42E-07 | >0.05 | >0.05 | OWBQRHWTHXHYCE |
| 10 | 1.28 | 365 | Steroids and steroidal derivative | (2R,3R,4S,5S,6R)-2-[(2S)-4-[(1R,2S,4S,6R,7S,8R,9S,12S,13S,16S,18R)- 16-[(2R,3R,4S,5S,6R)-4,5-dihydroxy-6-(hydroxymethyl)-3-[(2S,3R,4S,5R)-3,4,5-trihydroxyoxan-2-yl]oxyoxan-2-yl]oxy-6-methoxy-7,9,13-trimethyl-5-oxapentacyclo[10.8.0.02,9.04,8.013,18]icosan-6-yl]-2-methylbutoxy]-6-(hydroxymethyl)oxane-3,4,5-triol | 6.61E-08 | >0.05 | 0.00171061 | QAAUZCZXDMBHSC |
| 11 | 1.19 | 391 | Steroids and steroidal derivative | Methylprotogracillin | 1.75E-07 | >0.05 | 0.00038982 | LOSNTJHBTWBJCC |
| 12 | 1.55 | 400 | Steroids and steroidal derivative | (1R,2S,4S,6R,7S,8R,9S,12S,13R,16S)-16- [(2R,3R,4S,5S,6R)-3,4-dihydroxy-6-methyl-5-[(2S,3R,4S,5S,6R)-3,4,5-trihydroxy-6-(hydroxymethyl)oxan-2-yl]oxyoxan-2-yl]oxy-6-hydroxy-7,9,13-trimethyl-6-[(3R)-3-methyl-4-[(2S,3R,4S,5S,6R)-3,4,5-trihydroxy-6-(hydroxymethyl)oxan-2-yl]oxybutyl]-5-oxapentacyclo[10.8.0.02,9.04,8.013,18]icos-18-en-10-one | 4.31E-13 | >0.05 | 1.95E-06 | YEPLHFBCHYIXDM |
| 13 | 1.10 | 415 | Steroids and steroidal derivative | [(25R)-12-Oxo-5alpha-spirostane- 3beta,6alpha-diyl]bis(beta-D-glucopyranoside) | 3.82E-07 | >0.05 | >0.05 | HRNKURGIDWWIHJ |
| 14 | 1.18 | 471 | homoisoflavonoids | (3S)-5,7-dihydroxy-3-[(4-hydroxyphenyl)methyl]-2,3-dihydrochromen-4-one | 0.00055405 | >0.05 | >0.05 | FIASLUPJXGTCKM |
| 15 | 1.42 | 509 | Steroids and steroidal derivative | (1R,2S,3R,4R,5'R,6R,7S,8R,9S,12S,13R,14R,16R)- 5',7,9,13-tetramethylspiro[5-oxapentacyclo[10.8.0.02,9.04,8.013,18]icos-18-ene-6,2'-oxane]-3,14,16-triol | 1.11E-05 | 4.06E-14 | >0.05 | BSUPFYRQXCQGLJ |
| 16 | 1.43 | 511 | Steroids and steroidal derivative | 3beta-[4-O-(alpha-L-Rhamnopyranosyl)- beta-D-glucopyranosyloxy]spirosta-5,25(27)-diene-1beta-ol | 1.00E-18 | >0.05 | >0.05 | PFBUFKFWZHQYKN |
| 17 | 1.76 | 539 | Steroids and steroidal derivative | (1R,2S,3R,4S,6R,7R,8R)-1,2-dimethyl-8-propan-2-yltetracyclo[4.4.0.02,4.03,7]decane | 7.84E-05 | 7.46E-12 | >0.05 | XBWACJDEQIZTPR |
| 18 | 1.32 | 554 | homoisofomolavonoids | Methylophiopogonone A | 4.01E-07 | 1.26E-08 | >0.05 | MPUAHKMJGSMMIL |
| 19 | 1.28 | 556 | homoisoflavonoids | (3S)-5-hydroxy-3-[(4-hydroxy-3- methoxyphenyl)methyl]-7-methoxy-2,3-dihydrochromen-4-one | 0.000151668 | >0.05 | >0.05 | VSUZMLIEPGKFQG |
| 20 | 1.46 | 587 | homoisoflavonoids | Shancigusin F | 1.43E-08 | 4.69E-09 | >0.05 | HOPMWZPRWYIPRP |
| 21 | 1.54 | 590 | homoisoflavonoids | Analogue of Methylophiopogonanone A | 2.70E-11 | >0.05 | >0.05 | BXTNNJIQILYHJB |
| 22 | 1.79 | 603 | homoisoflavonoids | Methylophiopogonanone B | 0.001073072 | 1.98E-15 | >0.05 | UFMAZRUMVFVHLY |
| 23 | 1.41 | 614 | homoisoflavonoids | (+)-Syringaresinol | 6.35E-20 | >0.05 | >0.05 | KOWMJRJXZMEZLD |
| 24 | 1.54 | 620 | homoisoflavonoids | (3R)-3-[(3,4-dihydroxyphenyl)methyl]-5-hydroxy-7,8-dimethoxy-6-methyl-2,3-dihydrochromen-4-one | 1.29E-11 | 0.002270816 | >0.05 | JOUUNPLVSDYEPN |
| 25 | 1.37 | 626 | homoisoflavonoids | 4-[(3S,3aR,6R,6aR)-3-(1,3-benzodioxol-4-yl) -1,3,3a,4,6,6a-hexahydrofuro[3,4-c]furan-6-yl]-1,3-benzodioxole | 1.48E-07 | 4.44E-10 | >0.05 | MHWKVSWLLYGYHT |
| 26 | 1.47 | 645 | homoisoflavonoids | 5-Hydroxy-7-(4-hydroxy-3-methoxyphenyl)- 1-(4-hydroxyphenyl)hepta-4,6-dien-3-one | 1.46E-12 | 3.93E-09 | >0.05 | ZNVIPQYJPLZSBC |
| 27 | 1.50 | 648 | homoisoflavonoids | (2R,3R)-3-(1,3-benzodioxol-5-ylmethyl)- 2,5,7-trihydroxy-6,8-dimethyl-2,3-dihydrochromen-4-one | 1.65E-10 | 1.83E-08 | >0.05 | HUGFWFLIFYXRSJ |
| 28 | 1.36 | 657 | homoisoflavonoids | (3R)-5,7-dihydroxy-3-[(4-hydroxy- 3-methoxyphenyl)methyl]-6,8-dimethyl-2,3-dihydrochromen-4-one $ (3S)-5,7-dihydroxy-3-[(4-hydroxy-3-methoxyphenyl)methyl]-6,8-dimethyl-2,3-dihydrochromen-4-one | 7.29E-17 | 3.31E-06 | >0.05 | DPWJUIYOIZDQPQ |
| 29 | 1.48 | 664 | homoisoflavonoids | (3r)-3-(1,3-Benzodioxol-5-ylmethyl)-2,3- dihydro-7-hydroxy-5-methoxy-6,8-dimethyl-4h-chromen-4-one | 4.39E-13 | 3.72E-10 | >0.05 | YCKJMWYVLFOHAG |
| 30 | 1.51 | 667 | Steroids and steroidal derivative | Analogue of Ophiopogonin B | 1.04E-17 | >0.05 | 0.04488626 | OWGURJWJHWYCIQ |
| 31 | 1.4187 | 678 | Steroids and steroidal derivative | Ophiopogonin D | 1.07E-16 | >0.05 | >0.05 | FHKHGNFKBPFJCB |
| 32 | 1.47 | 679 | homoisoflavonoids | Medioresinol | 1.11E-05 | 0.000630218 | >0.05 | VJOBNGRIBLNUKN |
| 33 | 1.66 | 686 | homoisoflavonoids | (3r)-2,3-Dihydro-7-hydroxy-5-methoxy-3-(4-methoxybenzyl)-6,8-dimethyl-4h-chromen-4-one | 0.000827124 | 1.01E-29 | >0.05 | OXTVCCMEMSFLOW |
| 34 | 1.41 | 687 | Steroids and steroidal derivative | [(25R)-3,3-Dimethoxy-5alpha- spirostan-6alpha-yl]3-O-(beta-D-glucopyranosyl)-beta-D-glucopyranoside | 3.54E-18 | >0.05 | >0.05 | RNWZQKLXCVWMIK |
| 35 | 1.47 | 713 | homoisoflavonoids | OphiopogonanoneA | 9.99E-15 | 0.000330559 | >0.05 | QBRLTNYECODTFP |
| 36 | 1.49 | 717 | homoisoflavonoids | Guayarol | 2.58E-10 | 6.37E-09 | >0.05 | OIFFJDGSLVHPCW |
| 37 | 2.10 | 730 | Steroids and steroidal derivative | Ophiopogonin B | 1.07E-23 | >0.05 | >0.05 | OWGURJWJHWYCIQ |
| 38 | 1.84 | 737 | homoisoflavonoids | 2-Propenoic acid, 3-(4-hydroxy-3-methoxyphenyl)-, 2-(4-hydroxyphenyl)ethyl ester, (E)-;  4-Hydroxyphenethyl trans-ferulate | 0.002989818 | 3.50E-17 | >0.05 | JMSFLLZUCIXALN |
| 39 | 1.59 | 742 | homoisoflavonoids | Gibberellin A53 | 8.67E-11 | 6.56E-06 | >0.05 | CZEMYYICWZPENF |
| 40 | 1.27 | 750 | homoisoflavonoids | (2S,3S)-3-(1,3-benzodioxol-5-ylmethyl)- 2,5,7-trihydroxy-6,8-dimethyl-2,3-dihydrochromen-4-one | 0.029559454 | 2.08E-06 | >0.05 | HUGFWFLIFYXRSJ |
| 41 | 1.43 | 757 | homoisoflavonoids | 5,7-Dihydroxy-3-[(4-methoxyphenyl)methyl]- 8-methyl-4-oxochromene-6-carbaldehyde | 2.32E-13 | 5.75E-17 | >0.05 | MPUAHKMJGSMMIL |
| 42 | 1.77 | 759 | homoisoflavonoids | (2R,3R)-2,5,7-trihydroxy-3- [(4-methoxyphenyl)methyl]-6,8-dimethyl-2,3-dihydrochromen-4-one | 0.00335362 | 2.77E-28 | >0.05 | PLNGZQZXHIURST |
| 43 | 1.40 | 765 | homoisoflavonoids | Sesamin | 5.88E-12 | >0.05 | >0.05 | PEYUIKBAABKQKQ |
| 44 | 1.47 | 770 | Steroids and steroidal derivative | [(2S,3R,4R,5S,6S)-3,5-dihydroxy-2- [(2R,3R,4S,5S,6R)-5-hydroxy-2-[(1S,2S,4S,5'S,6R,7S,8R,9S,12S,13R,14R,16R)-16-hydroxy-5',7,9,13-tetramethylspiro[5-oxapentacyclo[10.8.0.02,9.04,8.013,18]icos-18-ene-6,2'-oxane]-14-yl]oxy-6-methyl-4-[(2S,3R,4S,5R)-3,4,5-trihydroxyoxan-2-yl]oxyoxan-3-yl]oxy-6-methyloxan-4-yl] acetate | 5.61E-14 | >0.05 | >0.05 | PYGKBFAELBHVIX |
| 45 | 1.54 | 780 | homoisoflavonoids | RubiadinOphiopogonanone F | 3.22E-11 | 0.002155776 | >0.05 | VYQRDDWHTRSYGE |
| 46 | 1.48 | 781 | homoisoflavonoids | Methylophiopogonanone A | 3.79E-17 | 2.68E-09 | >0.05 | BXTNNJIQILYHJB |
| 47 | 1.41 | 785 | homoisoflavonoids | Methylophiopogonone B | 1.48E-25 | >0.05 | >0.05 | BUTFXZVBLOLETI |
| 48 | 1.76 | 790 | homoisoflavonoids | Dihydrodesmethoxycurcumine a | 0.005197099 | 3.65E-27 | >0.05 | ZNVIPQYJPLZSBC |
| 49 | 1.57 | 802 | homoisoflavonoids | (3r)-2,3-Dihydro-7-hydroxy-5-methoxy-3-(4-methoxybenzyl)-6,8-dimethyl-4h-chromen-4-one | 1.76E-09 | 1.16E-05 | >0.05 | OXTVCCMEMSFLOW |
| 50 | 1.61 | 809 | homoisoflavonoids | Methylophiopogonanone B | 2.70E-06 | 2.57E-27 | >0.05 | UFMAZRUMVFVHLY |
| 51 | 1.04 | 820 | Steroids and steroidal derivative | (22S)-1beta-[2-O-(alpha-L-Rhamnopyranosyl)- 3-O-(beta-D-xylopyranosyl)-beta-D-xylopyranosyloxy]-16beta-(beta-D-glucopyranosyloxy)cholesta-5-ene-3beta,22-diol | >0.05 | 0.046656253 | >0.05 | IUYQQZFQIDNSNK |
| 52 | 1.54 | 827 | homoisoflavonoids | (3R,4R)-3,4-bis(1,3-benzodioxol-5- ylmethyl)-3-hydroxyoxolan-2-one | 0.010170089 | 3.88E-25 | >0.05 | OTWLSQPCSOEBAY |
| 53 | 1.06 | 841 | Steroids and steroidal derivative | (25s)-spirost-5-ene-1beta,3beta-diol 1- O-beta-d-fucopyranoside | 3.81E-08 | >0.05 | >0.05 | KIAIAZNJHWWAFM |
| 54 | 1.99 | 848 | homoisoflavonoids | (3r)-3-(1,3-Benzodioxol-5-ylmethyl)-2,3-dihydro- 7-hydroxy-5-methoxy-6,8-dimethyl-4h-chromen-4-one | 0.018354366 | 3.96E-26 | >0.05 | YCKJMWYVLFOHAG |
| 55 | 1.93 | 858 | Steroids and steroidal derivative | (1R,2S,3R,4R,5'R,6R,7S,8R,9S,12S,13R,14R,16R)-5',7,9,13- tetramethylspiro[5-oxapentacyclo[10.8.0.02,9.04,8.013,18]icos-18-ene-6,2'-oxane]-3,14,16-triol | >0.05 | 4.10E-09 | >0.05 | BSUPFYRQXCQGLJ |
| 56 | 1.50 | 892 | homoisoflavonoids | 5,7-Dihydroxy-3-[(4-methoxyphenyl) methyl]-8-methyl-4-oxochromene-6-carbaldehyde | 0.013280338 | 5.43E-22 | >0.05 | MPUAHKMJGSMMIL |
| 57 | 1.68 | 918 | fatty acid | 1-Pent-2-enylcyclohexa-1,3-diene | 7.01E-12 | 0.003001245 | >0.05 | AIELHXXAVIMENK |
| 58 | 1.63 | 919 | fatty acid | beta-Aromadendrene | 1.75E-11 | 0.000957716 | >0.05 | ITYNGVSTWVVPIC |
